# A mosaic gastruloid model highlights the developmental stage-specific restriction of cell competition in mammalian pre-gastrulation

**DOI:** 10.1101/2025.01.25.634859

**Authors:** J.D. Frenster, S. Babin, J.B. Josende Garcia, P. Casani Galdon, P. Pascual-Mas, G. Robertson, J. Garcia Ojalvo, A. Martinez Arias

## Abstract

In early mammalian embryogenesis, the selective elimination of suboptimal cells is critical for developmental integrity. Cell competition (CC) is a cell non-autonomous quality control in which “winner” cells eliminate viable but suboptimal “loser” cells based on their relative difference in fitness. Due to its central role in fitness perception, loss of p53 results in the emergence of “supercompetitor” cells, which stand at the apex of cell competition and induce apoptosis in neighboring wild-type (WT) cells.

Here, we investigate CC dynamics using mosaic 3D mouse gastruloids, an embryonic stem cell (ESC)-based *in vitro* model of gastrulation, composed of defined numbers of WT and p53-KO cells. In mosaic gastruloids, even low numbers of p53-KO cells robustly outcompete WT cells, and introduction of as few as two p53-KO cells is sufficient to measurably impair neighboring WT cell growth. CC in gastruloids is independent of cell proliferation rates, nutrient availability, or reactive oxygen species (ROS), and not influenced by Nodal and ERK signaling. However, we observe that Wnt and BMP signaling protect from CC, which is exclusively driven by intrinsic apoptosis, as indicated by Bcl2-mediated complete rescue of WT cells. During gastruloid development, CC is temporally restricted to a window of 48–96 hours after aggregation, mirroring embryonic days E5.5–E7.5 in the mouse. Heterochronic mosaic gastruloid experiments demonstrate that relative differences in pluripotency levels are neither necessary nor sufficient to cause supercompetition, but that CC is contingent on both competitors residing within the developmental window permissive to CC. Neither pluripotent mosaic 3D aggregates, nor 3D EpiGastruloids, which model more advanced developmental processes, display any competition, supporting the hypothesis that developmental CC is specific to the onset of gastrulation. Our findings offer insights into the mechanisms of cell fitness evaluation in mammalian embryogenesis and establish gastruloids as a powerful 3D model for investigating developmental stage-specific cell competition.

## Introduction

During mammalian development a single totipotent zygote, gives rise to the pluripotent cells of the blastocyst inner cell mass, which go on to form the epiblast and engages its cells in the process of gastrulation: a stereotyped series of cell movements associated with fate specification that lay down the body plan ^1^. Gastrulation is a central and conserved process in the development of all animals that involves organized cell fate assignments and stereotyped cell movements. In mammals it occurs against a background of rapid cell proliferation which increase by two orders of magnitude the 300/400 cell population of the epiblast ^2^.

In the mouse embryo, gastrulation starts at about E6.5 and, during the 24 hours preceding this process, there is a wave of apoptosis in the epiblast ^3, 4^ that eliminates up to 35% of the cells ^5^. While some of this apoptosis reflects autonomous death of severely compromised cells, it is complemented by a cell non-autonomous mechanism which ensures that among the viable cells only the fittest prevail. It has been suggested that this event reflects a process of *cell competition* (CC) in which cells compare their relative developmental fitness to that of their neighbors and less fit cells are eliminated. Surviving “fitter” cells are designated as “winner” cells, while their relatively less fit neighbors become “loser” cells and undergo apoptosis ^6-8^.

This relative nature of cell competition was first identified in *Drosophila*, where *Minute* mutant cells are outcompeted by wild-type (WT) cells ^9^, while healthy WT cells in turn can be induced to undergo apoptosis when exposed to so called “supercompetitor” cells overexpressing dMYC ^10^. In mammalian development, loser cells can be defined by a number of non-lethal defects, such as low cMyc expression ^11-14^, impaired Hippo ^15^ or BMP signaling ^5^, autophagy deficiencies ^12^, mitochondrial defects ^16^, or aneuploidy ^12^. Loser cell fate has been reported to lead to p53 protein stabilization which results in subsequent apoptosis ^5, 14, 17^. Due to its central role in fitness perception, loss of the p53 tumor suppressor results in the emergence of supercompetitor cells, which stand at the apex of cell competition and cause apoptosis in healthy neighboring wild-type (WT) cells ^18^. Besides defects and stress responses, relative differences in pluripotency levels have also been proposed to be a factor in cell competition, as cells of more naïve pluripotency outcompete less pluripotent cells after blastocyst co-injection ^19^, which fits to the positive correlation of pluripotency level and cMyc expression ^13^.

Despite the existence of many examples of intrinsic cellular variables associated with relative fitness levels, little is known about the nature of the dynamic crosstalk between competing cells. While cell competition in 2D epithelia may to a large part be driven by mechanical crowding and extrusion ^17, 20^, the mechanism facilitating communication and execution of cell competition in 3D tissues remain unclear. Additionally, the temporal regulation and control of cell competition is incompletely understood. These gaps in our understanding are in part due to the limited accessibility of embryos and the biological limitations of 2D models.

Here, we use mouse gastruloids as a novel platform to study mammalian cell competition during embryogenesis. Gastruloids are a 3D pluripotent stem cell-based *in vitro* model of mammalian perigastrulation development. Exploiting the robustness, reproducibility, and high throughput of the system we make a number of quantitative observations that validate them as a tool to study cell competition in mammalian development and that reveal new features of the process. We find that introducing a small number of p53-knockout (p53KO) supercompetitor cells into WT gastruloids suffices to induce intrinsic apoptosis in neighboring WT cells as a response to non-autonomous cell competition. Unlike in previous 2D systems, cell competition in gastruloids is independent of nutrient availability, total cell numbers, or mechanical compression, but instead is regulated by developmental signals. The tightly controlled temporal dynamics of cell competition in gastruloids mirror the events in the embryo and highlight the existence of a developmentally restricted window of time permissive for cell competition.

## Results

### 2D cell competition occurs only at confluence during differentiation

To investigate cell competition in a mammalian developmental system, we generated multiple fluorescently distinct mouse embryonic stem cell (ESC) clones from the same parental E14Tg2A line. We use an H2B-mCherry labeled wild-type (WT) clone as the primary “cell-under-investigation” (green, “WT-mCherry”) and an H2B-emiRFP670 labeled WT clone as the competitive neighborhood environment (magenta, “WT-emiRFP”) (**Extended Data Fig. 1a**). Cell competition is believed to be based on a cell’s intrinsic fitness perception and a comparison of it to its neighbors’ fitness. Given that the p53 machinery functions as a central stress response hub, with elevated p53 protein levels indicating higher levels of cellular stress, *Trp53*-knockout (hereafter: p53KO) cells are expected to perceive themselves as ‘flawless’ and thereby at the apex of cell competition (“supercompetitors”). In agreement with this, p53KO cells have previously been demonstrated to behave as supercompetitors in 2D differentiation settings ^5, 18^. Building on these observations and to create a system of unequal competitive strength, we generated a p53KO daughter clone from the WT-emiRFP clone by CRISPR-Cas9 mediated +1 frameshift in exon 4 of the *Trp53* gene (**Fig. 1a; Extended Data Fig. 1a-d**) (magenta, “p53KO-emiRFP”). This p53 loss-of-function did not alter colony morphology or alkaline phosphatase activity in maintenance culture (FBS+LIF) and did not affect doubling times relative to the parental WT-emiRFP clone (**Fig. 1b; Extended Data Fig.1a**).

**Figure 1.**
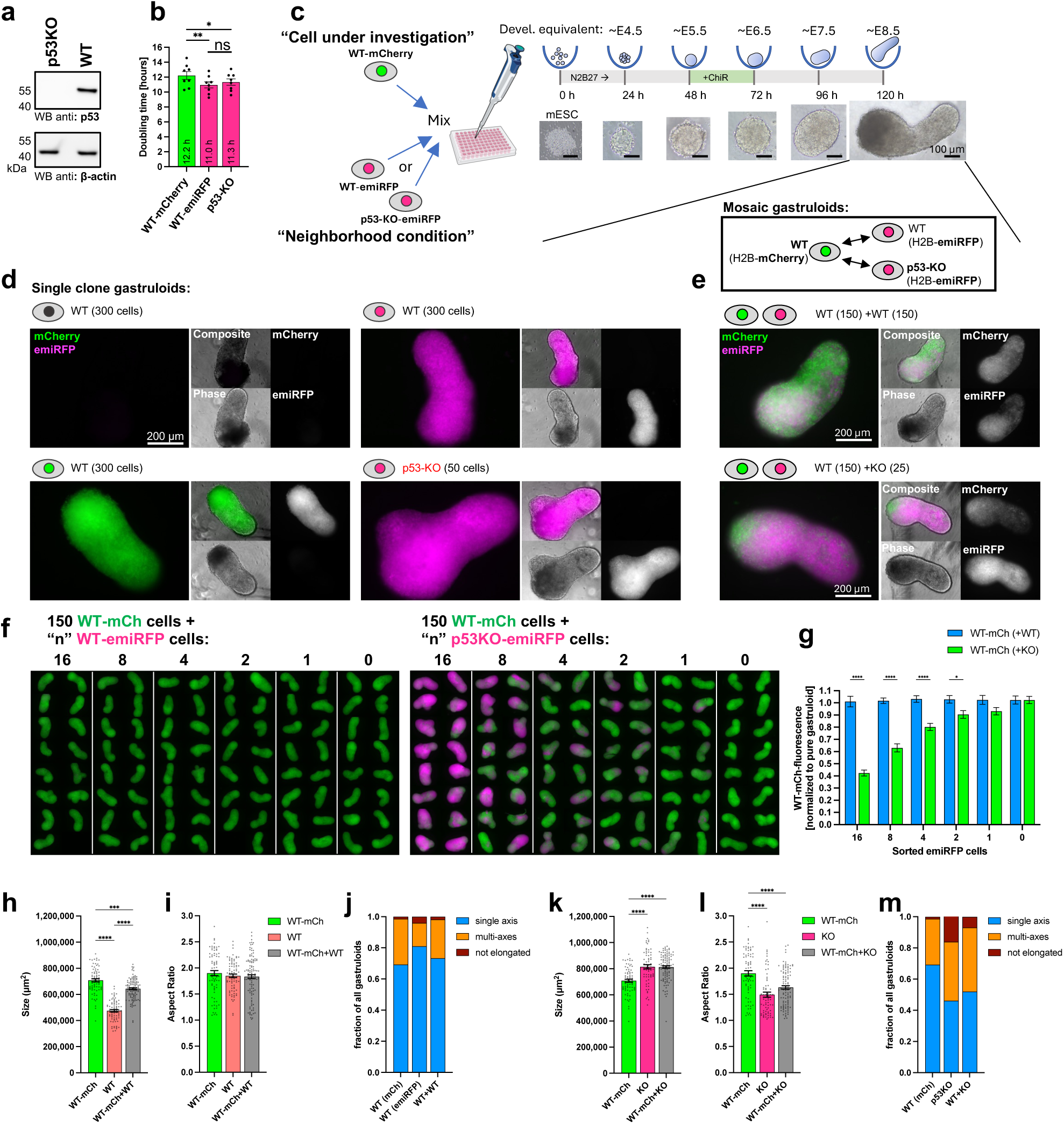
p53-knockout cells act as supercompetitors in mosaic 3D gastruloids. **a**, Western blot analysis of p53KO and WT cells stained against p53 (upper panel) and loading control β-actin (lower panel) confirms loss of p53 protein (Extended Data Fig. 1d). **b**, Doubling times of clones used in this study. n=8-9 independent experiments. *, p<0.05; **, p<0.01; ns, not significant. **c**, Schematic of mosaic gastruloid formation protocol. WT-mCherry mESCs are mixed with WT-emiRFP or p53KO-emiRFP mESCs and aggregated in U-bottom wells. Developmental equivalent timepoints of mouse development are annotated at the top and brightfield example images of gastruloids at respective timepoints are shown below. Scale bar denotes 100 µm. **d**, Representative widefield fluorescence microscopy images of gastruloids from parental WT E14 cell line (top left), WT-mCherry cells (bottom left), WT-emiRFP cells (top right), or p53KO-emiRFP cells (bottom right) at 120h. Scale bar denotes 200 µm. **e**, Representative widefield fluorescence microscopy images of mosaic gastruloids from 150 WT-mCherry and 150 WT-emiRFP cells (top panel) or 150 WT-mCherry and 25 p53KO-emiRFP cells (bottom panel) at 120h. Scale bar denotes 200 µm. **f**, Cell number titration of WT-emiRFP (left panel) or p53KO-emiRFP (right panel) cells seeded in mosaic gastruloids together with 150 WT-mCherry cells per gastruloid. Representative widefield fluorescence microscopy montage of 120h gastruloids depicted as overlay of mCherry and emiRFP channels. Numbers of emiRFP cells seeded per mosaic gastruloid are denoted over each panel. Seeding by fluorescence activated cell sorting in single cell purity mode. **g**, Quantification of panel **f**. mCherry fluorescent intensity per gastruloid normalized to pure mCherry gastruloids (0) as a measure of WT-mCherry cell growth. n=48 gastruloids per condition from 3 independent experiments. **h-m**, Morphological description of monoclonal and mosaic gastruloids depicting size quantified as 2D area of widefield images (**h,k**), aspect ratio of long axis to short axis of gastruloids (**i,l**), and quantification of single vs multi-axes formation (**j,m**). n=68-111 gastruloids per condition. ***, p<0.001; ****, p<0.0001. (Single and multi-axis examples and analysis mask depicted in Extended Data Fig. 2c). All values depicted as mean ±SEM throughout this figure.

We first assessed the clones’ competitive behavior in a 2D co-culture setting by mixing WT-mCherry cells with either WT-emiRFP or p53KO-emiRFP cells. In pluripotent co-passaging conditions, we observed a slow drift in favor of both emiRFP clones (**Extended Data** Fig. 1e), which was independent of p53 deletion status (**Extended Data Fig. 1f**). We then allowed cells to exit naïve pluripotency toward a primed state (N2B27+AA+FGF) without passaging. In these conditions, co-cultured WT-mCherry and WT-emiRFP cells displayed no difference in their growth behavior, while the presence of p53KO cells impaired the growth of co-cultured WT cells and resulted in a decline of WT cell numbers over time (**Extended Data Fig. 1g-h**), confirming previous studies on p53KO supercompetition (Bowling et al, 1018; Perez Montero et al., 2024). However, we observed that the temporal onset of this cell competition-like effect was dependent on the initial seeding numbers, and competition only became apparent when the sum of cells per well reached confluence, at which point the daily medium changes became insufficient to sustain the number of cells per well (**Extended Data Fig. 1h-i, o**). This same effect was not observed in pluripotent confluent cultures (**Extended Data** Fig. 1j-k). Similarly, when 2D co-cultures were allowed to exit naïve pluripotency while being passaged as Epi-like clumps every 72h to avoid overcrowding, they did not compete despite growing as mixed mosaic colonies (**Extended Data Fig. 1l-n**).

These observations show that, in adherent 2D differentiating conditions, p53KO-mediated cell competition only occurs once confluence is reached. This suggests that, in these conditions, cells either compete for nutritional resources or limited space, which may or may not reflect cell competition in development. We thus moved on to investigate cell competition in gastruloids, a 3D model of mammalian gastrulation that has been shown to more closely reflect the situation in the embryo ^21-23^, while maintaining a larger medium-to-cell-number ratio throughout the differentiation timeline.

### p53KO cells act as supercompetitors in mosaic 3D gastruloids

Gastruloids are formed by aggregation of a defined number of ESCs that over five days and under defined culture conditions develop into elongated and polarized structures that represent the embryo at E8.5 ^21-23^ (**Fig. 1c**). WT-mCherry, WT-emiRFP, and p53KO-emiRFP clones generate gastruloids in a manner comparable to their parental culture of origin (E14Tg2A) (**Fig. 1d**), with the difference that that p53KO gastruloids require fewer starting cells for form gastruloids (50) than WT ones.

Mixing 150+150 WT-mCherry and WT-emiRFP cells (WT+WT) during aggregation results in mosaic gastruloids composed of equal contribution of both WT clones (**Fig. 1e**). However, even mixing 150 WT-mCherry and 25 p53KO-emiRFP cells (WT+p53KO), results in mosaic gastruloids mostly composed of p53KO-emiRFP cells (**Fig. 1e**). This suggests an impairment of growth or survival of the WT-mCherry cells conditioned by the p53 status of their neighboring cells. To test the relation between cell stoichiometry and competitive outcome, we titrated the number of p53KO-emiRFP cells seeded together with WT-mCherry cells during aggregation and observed that as few as two starting p53KO-emiRFP cells per gastruloid suffice to measurably impair the growth of the co-aggregated 150 WT cells (**Fig. 1f-g**). Together, these data indicate that p53KO cells behave as supercompetitors in 3D gastruloids and that very few p53KO cells suffice to impair the growth or survival of neighboring WT cells.

Following these observations, we interrogated whether the competitive interactions had consequences for mosaic gastruloid morphology or cell fates. In monoclonal form, WT-mCherry gastruloids were slightly larger than WT-emiRFP gastruloids, but smaller than p53KO-emiRFP gastruloids (**Fig. 1h,k; Extended Data Fig. 2c**), and p53KO gastruloids displayed more morphological irregularities than WT gastruloids (**Fig. 1i-m**). Mosaic WT+WT gastruloids resulted in the average size of the two clones, without affecting aspect ratio or axis formation efficiency (**Fig. 1h-j**). In contrast, mosaic WT+p53KO gastruloids grew to the same size of monoclonal p53KO gastruloids, despite starting with only half the number of p53KO cells, suggesting compensatory growth during competitive interactions (**Fig. 1k-m**).

Knockout of *Trp53* and its homologues has previously been described to impair mesendoderm formation in mouse development ^24^. We thus set out to test how germ layer specification was influenced in our mosaic competitive conditions. While WT+WT gastruloids form organized regions from all three germ layers, WT+p53KO gastruloids at 120h display lower and more disorganized centers of FoxA2 and E-cadherin expression, suggesting impaired endoderm formation (**Extended Data Fig. 2a-b**). In contrast, patterned Tbx6 and Bra expression (mesoderm), was not impaired in WT+p53KO gastruloids. Transcriptomic analysis of fluorescence activated cell sorting (FACS) isolated cells from mosaic gastruloids confirm that p53KO cells in 120h gastruloids were biased towards paraxial mesoderm and caudal epiblast formation, while being impaired in hematoendothelial and PGC development (**Extended Data Fig. 2d-e**). Overall, p53KO cells in 120h mosaic gastruloids resembled more posterior compartments of the embryo when compared to WT gastruloids (**Extended Data Fig. 2f**). WT-mCherry cells from mosaic WT+p53KO gastruloids displayed only slight differences in their lineage marker expression profile from those in WT+WT mosaics, but nonetheless followed the same trend as p53KO cells, suggesting non-autonomous lineage formation effects due to the presence of p53KO cells. Surprisingly, no significantly differential expressed genes were detected between WT-mCherry cells from mosaic WT+WT or WT+p53KO gastruloids at 48h and 72h (**Extended Data Fig. 2g**), suggesting that the presence of p53KO cells does not influence the neighboring WT transcriptome until later stages.

### Cell competition is temporally but not spatially restricted in gastruloids

Gastruloids are spatially organized 3D structures with established body axes. We thus aimed to investigated whether competing clones undergo spatial sorting, and whether loser populations are preferentially abolished in specific regions of the gastruloid. In both, WT+WT as well as WT+p53KO gastruloids, cells from different clones remained evenly distributed throughout as detected by confocal microscopy, independent of their winner or loser status (**Fig. 2a-b**). Using 3D segmentation and neighborhood analysis, we observed no striking bias of any clone being preferentially surrounded by “self” vs “other” clones (**Fig. 2c-d**). Furthermore, we observed no preferential spatial distribution of any clone along the center-to-edge axis of gastruloids (**Fig. 2c-e**). Together, these observations suggest that cell competition and localization of different clones within gastruloids is not spatially restricted.

**Figure 2.**
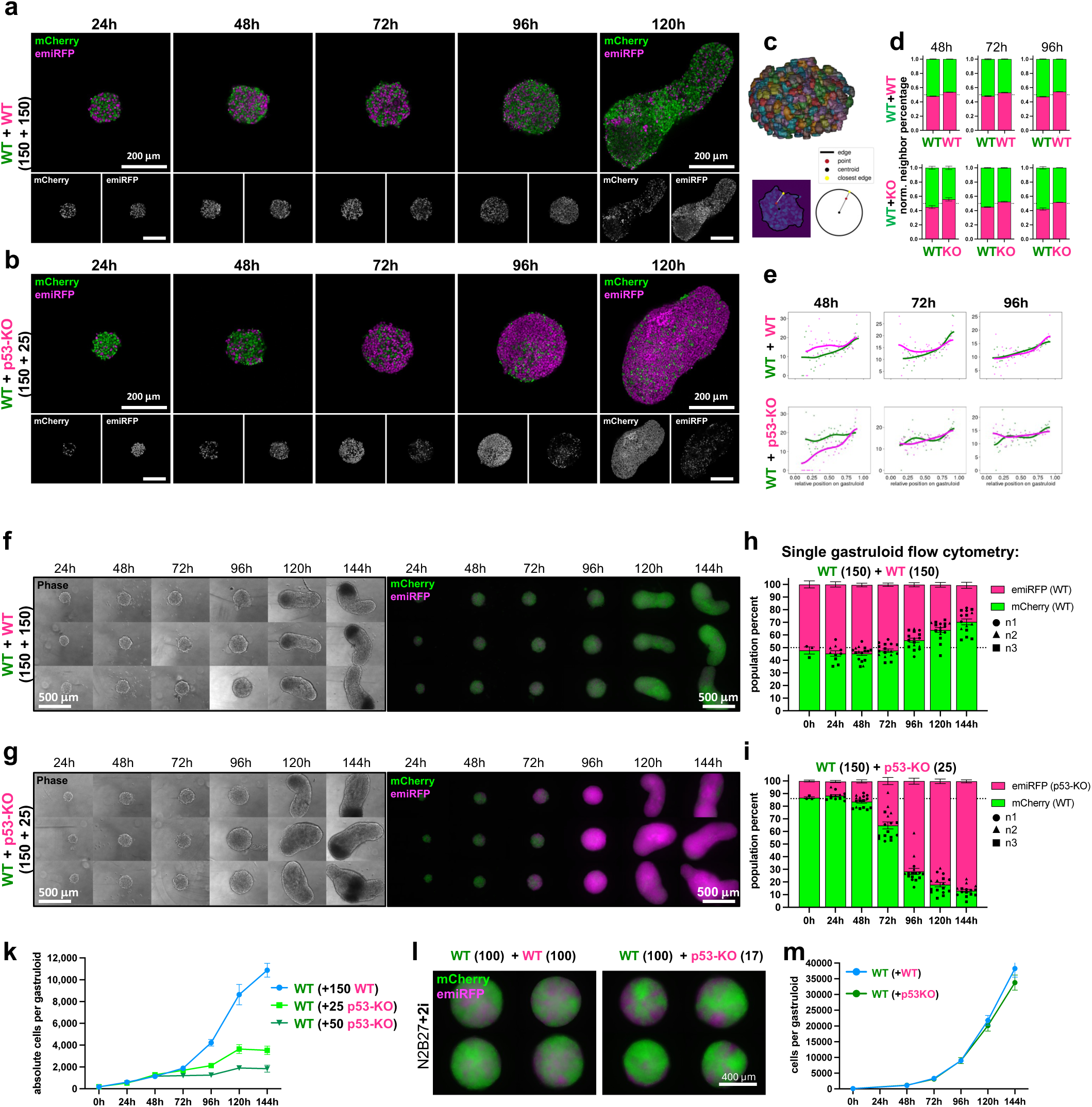
Cell competition is temporally but not spatially restricted in gastruloids. **a-b**, Confocal cross section images of mosaic gastruloids at different timepoints from 24-120h after aggregation. Representative images of WT-mCherry+WT-emiRFP gastruloids (**a**) and WT-mCherry+p53KO-emiRFP gastruloids (**b**) depicted. Scale bar denotes 200 µm. **c**, Example rendering of 3D segmented nuclei in 48h gastruloid (top) and schematic of center to edge analysis. **d**, Neighborhood analysis of 3D segmented mosaic gastruloids at 48h, 72h, 96h. Bar graphs depict the normalized percentage of mCherry (green) or emiRFP (magenta) neighbor cells amongst the nearest 20 nuclei centered around WT-mCherry, WT-emiRFP, or p53KO-emiRFP nuclei, as denoted below bars. Mean of all nuclei within 3-5 gastruloids per condition. **e**, Center-to-edge analysis of 3D segmented gastruloids. Depicts the normalized number of mCherry (green) or emiRFP (magenta) nuclei within an expanding shell from center (left) to edge (right) of mosaic gastruloids at 48h, 72h, or 96h. (n=3-5 gastruloids per condition) **f-g**, Widefield phase contrast and fluorescent images of 3 representative mosaic WT+WT (**f**) and WT+p53KO (**g**) gastruloids per timepoint from 24-144h after aggregation. Scale bar denotes 500 µm. **h-i**, Single gastruloid flow cytometry analysis depicting population composition of mosaic WT+WT (**h**) or WT+p53KO gastruloids (**i**), as percentage of mCherry+ (green) and emiRFP+ (magenta) cells. n=12-16 gastruloids from 3 independent experiments (24-144h), or n=3 (0h). Fitted curve in Extended Data Fig. 3b. **k**, Absolute growth curves of WT-mCherry cells in mosaic gastruloids seeded together with 150 WT-emiRFP (blue), 25 p53KO-emiRFP (light green), or 50 p53KO-emiRFP (dark green) cells (n=12 gastruloids per datapoint from 3 independent experiments). **l**, Representative widefield fluorescent images of pluripotent mosaic 3D aggregates grown in N2B27+2i. Mosaic aggregates of WT+WT (left) or WT+P53KO (right) gastruloids imaged at 120h after aggregation. Scale bar denotes 400 µm. **m**, Absolute growth curves of WT-mCherry cells in mosaic pluripotent aggregates seeded together with WT-emiRFP (blue) or p53KO-emiRFP (dark green) cells (n=12 gastruloids per datapoint from 3 independent experiments). All values depicted as mean ±SEM throughout this figure.

Gastruloids in many aspects resemble the same developmental time course of the mouse embryo. We therefore interrogated whether cell competition was uniform through time, or concentrated during specific developmental events. In mosaic gastruloids, p53KO-emiRFP cells became the dominant population around 72-96h after aggregation (**Fig. 2f-g**), despite their lower starting cell numbers (150 WT+25 p53KO cells). We conducted single gastruloid flowcytometry to investigate these temporal dynamics in a quantitative manner (**Extended Data Fig. 3a**). Population proportions in WT+WT mosaic gastruloids (50% + 50%) remain stable with a slight drift in favor of the WT-mCherry cells starting at 96-120h (**Fig. 2h**). In contrast, the proportions of WT cells in WT+p53KO mosaic gastruloids remain stable only until 48h (86% + 14%), after which the population percentages invert with the p53KO cells outcompeting the WT cells between 48-96h (**Fig. 2i; Extended Data Fig. 3b**). Mouse gastruloids at 48h, 72h, and 96h correspond to the embryonic epiblast around E5.5, E7.5, and the primitive streak around E8.0, respectively (**Fig. 6e**). The timing of cell competition in gastruloids thus corresponds to the apoptosis peak in mouse embryos ^3^.

Our analysis additionally revealed a high uptake of fluorophore-tagged nuclear material of the respective neighboring populations within mosaic gastruloids, likely due to phagocytosis (**Extended Data Fig. 3c-d**). At 72-96h after aggregation, up to 70% of H2B-emiRFP cells have taken up H2B-mCherry fluorescent material from neighboring cells. However, p53KO status did not influence this percentage, suggesting that phagocytosis might not be an active mechanism of cell competition in this system, but rather a mechanism of clearing debris (**Extended Data Fig. 3e**).

To understand whether the observed relative changes in population percentage (**Fig. 2h-i**) are caused by overgrowth of the p53KO cells, or decrease in the WT-mCherry cell numbers, we generated absolute growth curves of each clone population. This was achieved by transferring individual mosaic gastruloids to flat bottom plates, dissociating and fixing single cells without aspiration, followed by tile imaging and automated segmentation of fluorescent nuclei (**Extended Data Fig. 3f-g**). This method validated the previously observed relative population changes (**Extended Data Fig. 3h**) and revealed that absolute growth curves of WT-mCherry cells are impaired in mosaic gastruloids containing p53KO cells, but not when confronted by other WT cells (**Fig. 2k; Extended Data Fig. 3i-j).** This growth impairment is relative to the number of p53KO cells added and exacerbated when adding 50 instead of 25 p53KO cells into the mosaic gastruloids. Taken together, WT-mCherry cells are thus responsive to the p53 status of their neighbors, and the number of their p53KO neighbors.

As previously reported in 2D cultures ^12, 18^(**Extended Data Fig. 1f, j**), developmental cell competition does not take place in pluripotent cells. In accordance with this, WT and p53KO cells in 3D aggregates blocked from exiting pluripotency (2i, 2i+LIF) grow without impairment or competition, despite consisting of up to 70,000 cells and thus accounting for 3D physiological factors like nutrient scarcity and signaling molecule accumulation (**Fig. 2l-m; Extended Data Fig. 3k-n**).

Together, these data suggest that during gastruloids formation, the same WT-mCherry population can either co-develop with other WT cells by equally contributing to the final gastruloid or become impaired in their growth and/or viability when exposed to p53KO supercompetitor cells. This neighbor-dependent change in behavior rapidly manifests between 48h-96h after aggregation, which developmentally corresponds to the mouse embryo at E5.5-E7.5.

### Cell competition in gastruloids is mediated by intrinsic apoptosis

WT-mCherry cells exhibit near-exponential growth in WT+WT mosaic gastruloids, while their numbers stagnate after 48h in the presence of p53KO supercompetitor cells (**Fig. 2k**). To investigate whether this stagnation was caused by altered proliferation, we quantified the percentage of mitotic cells per cell population by phospho-Histone3 (pH3) staining and flowcytometry (**Fig. 3a; Extended Data Fig. 4a**). Surprisingly, WT-mCherry cells displayed the same percentage of mitotic cells independent of whether they were in mosaic gastruloids with other WT, or p53KO cells (**Fig. 3b**). Similarly, p53KO cells did not have an elevated percentage of mitotic cells compared to WT cells (**Extended Data Fig. 4b**). These data indicate that the observed differences in growth curves are not due to altered proliferation rates.

**Figure 3.**
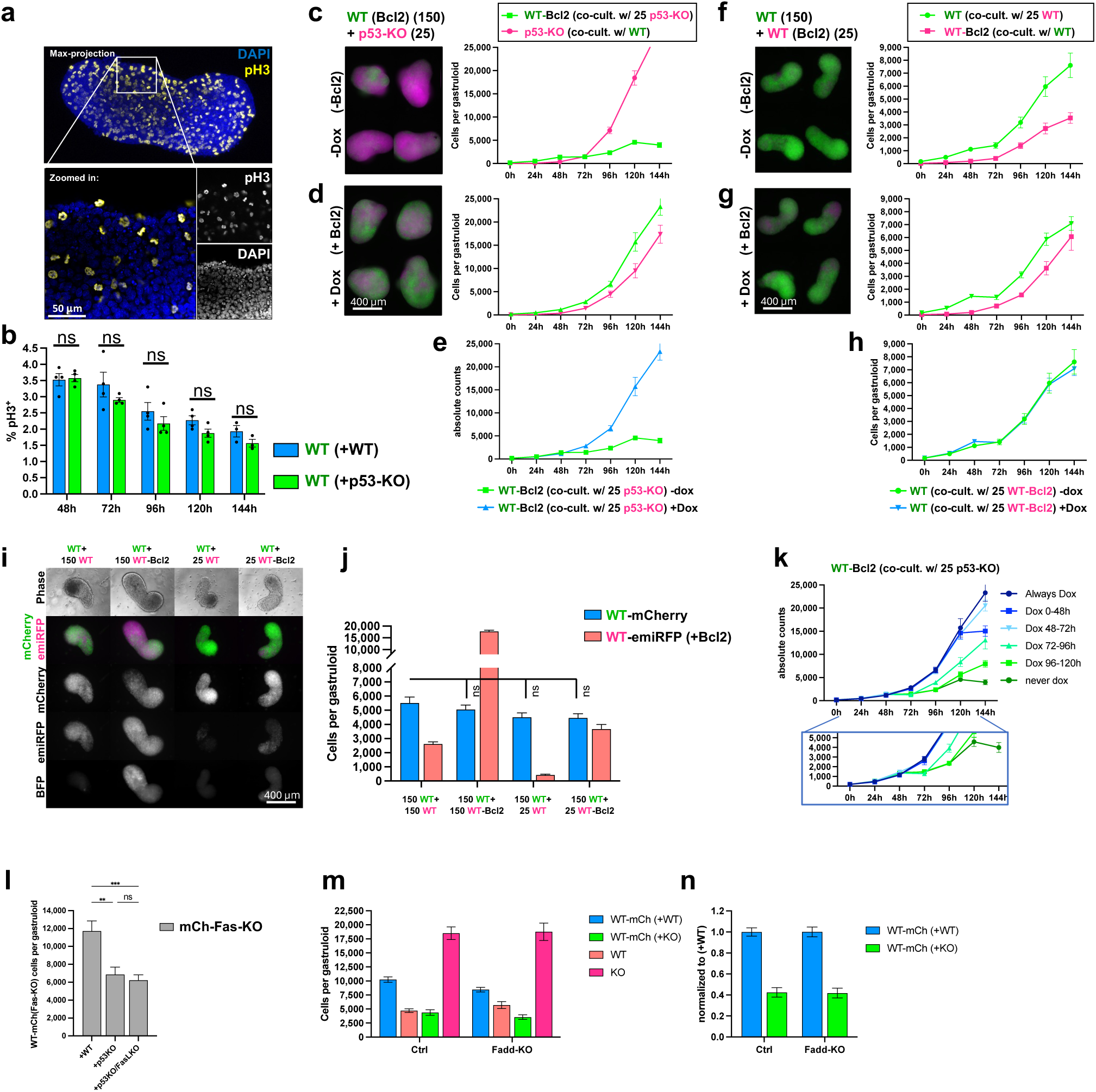
Cell competition in gastruloids is mediated by intrinsic apoptosis. **a**, Immunostaining micrograph of phospho-histone3 (pH3) stained gastruloid at 120h. Maximum intensity projection (top panel) and zoomed-in single confocal slices (lower panel) depicted as composite of DAPI (blue) and pH3 (yellow) channel. Scale bar denotes 50 µm. **b**, Flow cytometric quantification of phospho-histone3 positive percentage of WT-mCherry cells within WT+WT or WT+p53KO mosaic gastruloids at varying timepoints. n=4 independent experiments. **c-d**, Representative fluorescent widefield micrographs (left) of mosaic gastruloids at 120h containing p53KO-emiRFP cells and WT-mCherry cells with inducible Bcl2 expression cassette in the absence (**c**) or presence (**d**) of doxycycline induction, and corresponding quantified absolute growth curves (right panels). n=9 gastruloids from 3 independent experiments. **e**, Overlaid growth curves of WT-mCherry-Bcl2 cells from panels **c** and **d**, without and with doxycycline respectively. **f-g**, Representative fluorescent widefield micrographs (left) of mosaic gastruloids at 120h containing WT-mCherry cells and WT-emiRFP cells with inducible Bcl2 expression cassette in the absence (**f**) or presence (**g**) of doxycycline induction, and corresponding quantified absolute growth curves (right panels). n=9 gastruloids from 3 independent experiments. **h**, Overlaid growth curves of WT-mCherry cells from panels **f** and **g**, without and with doxycycline respectively. **i**, Mosaic gastruloids at 120h containing WT-mCherry cells together with WT-emiRFP or WT-emiRFP-Bcl2 cells. Scale bar denotes 400 µm. **j**, Quantification of absolute cell numbers of WT-mCherry, WT-emiRFP, or WT-emiRFP-Bcl2 cells from mosaic gastruloids at 120h as depicted in panel **i**. n=17-23 gastruloids from 4 independent experiments. **k**, Temporal Bcl2-overexpression series. Absolute growth curves of WT-mCherry-Bcl2 cells in mosaic gastruloids with p53KO-emiRFP cells, induced with doxycycline at varying time points as indicated. n=9 gastruloids from 3 independent experiments **l**, Absolute cell numbers of mCherry-Fas-knockout cells at 120h in mosaic gastruloids with WT-emiRFP, p53KO-emiRFP, or p53KO-FasLKO-emiRFP cells. n=12-18 gastruloids from 3 independent experiments. Statistical analysis between WT-mCherry counts across conditions depicted; ns, not significant. **m-n**, Absolute (**m**) or normalized (**n**) counts of clonal populations in mosaic gastruloids at 120h in control conditions, or after *Fadd*-knockout in mCherry cells. All values depicted as mean ±SEM throughout this figure.

To test whether cell competition in gastruloids is mediated by apoptosis of the WT cells, we generated a daughter clone of the WT-mCherry population carrying a doxycycline-inducible *Bcl2* overexpression construct. Bcl2 is a dominant inhibitor of apoptosis downstream of p53, Puma, and Noxa ^25^ (**Extended Data Fig. 4c-e**). In the absence of doxycycline, this WT-mCherry-Bcl2 clone was outcompeted by p53KO cells comparable to its parental clone (**Fig. 3c; Extended Data Fig. 4f**). However, upon doxycycline administration and Bcl2 expression, the WT-mCherry-Bcl2 clone remained unaffected by p53KO neighbor cells (**Fig. 3d-e, Extended Data Fig. 4f**), which mirrored recent observations in 2D supercompetition settings (Perez-Montero et al., 2024). Interestingly, the p53KO cells displayed impaired growth upon blocking of apoptosis in neighboring WT cells (**Extended Data Fig. 4g**), supporting the notion of compensatory growth upon competition. Taken together, inhibition of apoptosis by Bcl2 expression abolishes cell competition in gastruloids, corroborating that cell competition is executed through apoptosis of the ‘loser’ cells. p53KO (winner) cells are intrinsically resistant to stress-related apoptosis. It was thus paramount to test whether absence of apoptosis in one of the competing cell populations was sufficient to cause or mimic the cell competition effects we observed. We created an inducible Bcl2 expressing daughter clone of the WT-emiRFP cells and tested whether overexpression of Bcl2 and thus inhibition of apoptosis was sufficient to impair growth in the neighboring WT-mCherry clone (**Extended Data Fig. 4d-e**). Surprisingly, the WT-mCherry cells in mosaic gastruloids were unaffected by the overexpression of Bcl2 in neighboring WT-emiRFP cells (**Fig. 3f-h; Extended Data Fig. 4h**). Furthermore, seeding mosaic gastruloids with 25 or 150 cells of the WT-emiRFP or Bcl2-emiRFP clone, which resulted in a range of 400 -17,000 emiRFP cells per mosaic gastruloid at 120h, did not significantly alter the number of neighboring WT-mCherry cells in 120h gastruloids (**Fig. 3i-j**). This indicates that in WT+WT mosaic gastruloids, WT-mCherry cells are entirely unresponsive to the number of cells from their neighboring clones as long as there is no difference in competitive fitness. Bcl2 overexpressing apoptosis-deficient cells are unable to mimic the cell competition pressure exerted by the p53KO cells, indicating that the p53KO-mediated cell competition pressure is more than simply the absence of apoptosis in one of the competing populations.

Next, we made use of the inducible nature of the WT-*mCherry-Bcl2* expression clone in WT+p53KO mosaics, to test when during gastruloid formation cell competition induces apoptosis (**Extended Data Fig. 4i**). Continuous expression of *Bcl2*, or pulse induction of *Bcl2* expression after 48h of gastruloid formation, resulted in identical growth curves, indicating that the time of 0-48h of mosaic gastruloid formation, and thus the transition stage from pluripotency to a formative state, is irrelevant to cell competition (**Fig. 3k**). Induction of *Bcl2* after 72h rescued cell competition partially, and induction of *Bcl2* after 96h did not rescue WT cells from cell competition (**Fig. 3k**). These observations further support the notion that cell competition is restricted to the developmental stages modeled in 48-96h gastruloids, corresponding to the E5.5-E7.5 mouse epiblast.

Apoptosis can be initiated by intrinsic stress response or by extrinsic cell death signals. Cell competition has been reported to be executed via extrinsic apoptosis in hair follicle stem cell niche ^26^ but reported to be intrinsic in the embryo as determined by low levels of caspase 8 activation ^11^. Extrinsic apoptosis has classically been studied in the immune system, where natural killer (NK) cells and cytotoxic T-lymphocytes induce apoptosis in targeted cells via signaling through Fas ligand (*Fasl*) to Fas receptor (*Fas*) or tumor necrosis factor α (*Tnf*) to TNF receptor (*Tnfrsf1a*). RNAseq time course data from sorted mosaic gastruloids revealed that expression of both Fas receptor and TNF receptor is upregulated around 72-96h of gastruloid formation, overlapping with the time of cell competition (**Extended Data Fig. 4j**), but that only Fas ligand is upregulated concurrently, while *Tnf* expression peaked later. Intriguingly, p53KO cells expressed almost no Fas receptor, but higher levels of Fas ligand then their WT neighbors. We therefore created a Fas receptor knockout (Fas-KO) WT-mCherry daughter clone and a *Fasl* mutant (FasL-KO) p53KO-emiRFP daughter clone (**Extended Data Fig. 4k**) and tested whether absence of either receptor, ligand, or both would interrupt cell competition. Fas-KO-mCherry cells experienced cell competition stress, independent of whether they were in mosaic gastruloids with p53KO-emiRFP, or FasL-KO-p53KO-emiRFP supercompetitors (**Fig. 3l**), suggesting that the Fas ligand-receptor signaling pathway is not involved in cell competition. Extrinsic apoptosis signals from several TNF superfamily receptors get relayed via the Fas associated death domain (*Fadd*) in the signal receiving cells ^27^. However, even loss of this critical adaptor did not impair cell competition in Fadd-KO-mCherry cells compared to WT-mCherry cells (**Fig. 3m-n; Extended Data Fig. 4k**). Together, these data indicate that cell competition in gastruloids is strictly mediated by intrinsic apoptosis, and not a result of extrinsic apoptosis signaling.

Following the insight that cell competition in gastruloids is mediated by intrinsic apoptosis, we conducted 3D segmentation of Caspase 3 stained early, mid, or late apoptotic events in mosaic gastruloids and interrogated the cell identity of their neighborhood. Indeed, mid-apoptotic WT-mCherry cells had a higher probability of being surrounded by p53KO cells then by other WT cells at 72-96h (**Extended Data Fig. 5a-d**). Early apoptotic cells had a very slight bias toward the center of the gastruloid, particularly at 96h (**Extended Data Fig. 5e-f),** but this slight bias was insufficient to generate cell population gradients along the center to edge axis (**Fig. 2e**). Taken together, we find a trend of dying cells being preferentially surrounded by p53KO cells in more central areas of the gastruloid.

### Cell competition is not affected by gastruloid size, glucose availability, ROS, or cytoskeletal perturbations

We next tested what external variables might influence the apoptosis of loser cells. Loss of the tumor suppressor p53 has been linked to higher metabolic activity and a metabolic shift toward glycolysis (Warburg effect) ^28^, raising the possibility that p53KO-mediats competition could be driven by metabolic demand. Making use of the fact that the developmental processes in gastruloids scale with size within certain limits ^29, 30^, we seeded mosaic competition gastruloids with 200%, 50% and 25% of the normal starting cell number (600, 150, 75 total cell equivalents). As previously reported, the resulting gastruloid size and absolute cell numbers scaled linearly with the number of initially seeded cells (**Fig. 4a-b,d; Extended Data Fig. 6a-b).** However, the relative effect size of cell competition exerted by p53KO cells onto their neighbors, as measured by the ratio of WT-mCherry cells in WT+WT and WT+p53KO mosaic gastruloids, was unaffected by the eightfold difference in gastruloid size (**Fig. 4c**). Similarly, the temporal kinetics of cell competition centered around 72h after aggregation were unaffected by gastruloid size (**Fig. 4e**), standing in contrast to our observations of cell competition in 2D (**Extended Data Fig. 1h-i**). These data suggest that the onset of cell competition is determined by a developmental stage, and not a gastruloid size threshold.

**Figure 4.**
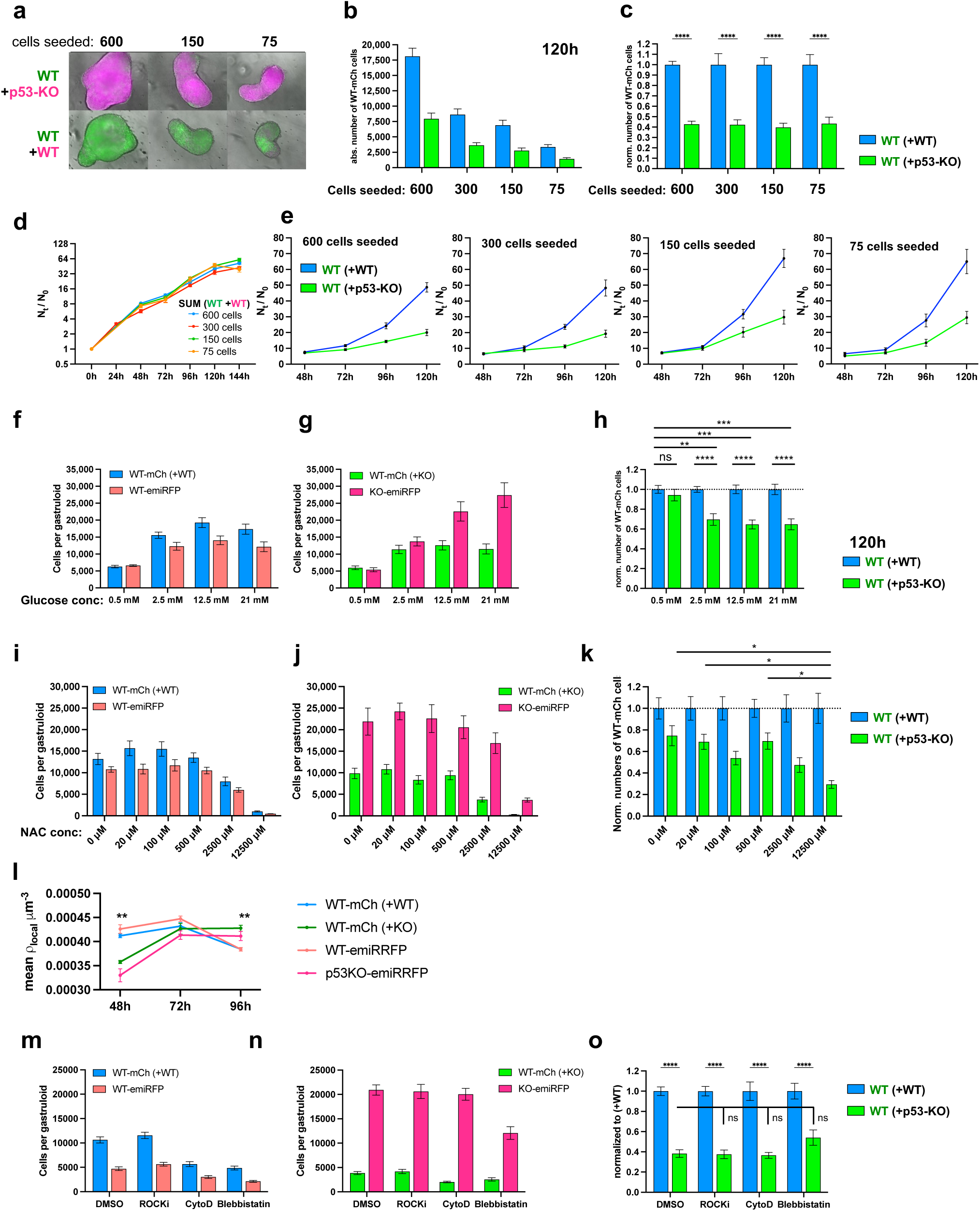
Cell competition is not affected by gastruloid size, glucose availability, ROS, or cytoskeletal perturbations. **a**, Representative fluorescent widefield micrographs of mosaic 120h gastruloids seeded from 600, 150, or 75 total cells. (corresponds to Extended Data Fig. 6a). **b**, Absolute counts of WT-mCherry cells at 120h in mosaic WT+WT (blue) or WT+p53KO (green) gastruloids. n=12 gastruloids from 3 independent experiments. **c**, Normalized counts of WT-mCherry cells at 120h in mosaic WT+WT (blue) or WT+p53KO (green) gastruloids. n=12 gastruloids from 3 independent experiments. Statistical analysis between WT+WT and WT+p53KO conditions within same cell seeding numbers depicted; ****, p<0.0001. **d**, Growth curves of sum of all cells within WT+WT gastruloids normalized to initial seeding numbers (N_t_/N_0_) demonstrates proportional scaling of growth behavior. n=12 gastruloids from 3 independent experiments each. **e**, Normalized growth curves of WT-mCherry cells in WT+WT (blue) and WT+p53KO (green) mosaic gastruloids from different initial seeding numbers demonstrate similar growth and competition kinetics. n=12 gastruloids from 3 independent experiments. **f-h**, Glucose and pyruvate limitation study. Absolute (**f-g**) and normalized (**h**) cell numbers for cell populations in WT+WT (**f**) or WT+p53KO (**g**) mosaic gastruloids at 120h, grown in varying concentrations of glucose. n=24 gastruloids from 4 independent experiments. Only statistical analysis between normalized WT-mCherry populations from WT+WT vs WT+p53KO gastruloids depicted; **, p<0.01; ***, p<0.001; ****, p<0.0001; ns, not significant. **i-k**, Reactive oxygen species (ROS) scavenger titration study. Absolute (**i-j**) and normalized (**k**) cell numbers for cell populations in WT+WT (**i**) or WT+p53KO (**j**) mosaic gastruloids at 120h, grown in varying concentrations of the ROS scavenger N-acetyl-L-cysteine. n=11 gastruloids from 3 independent experiments. Only statistical analysis between normalized WT-mCherry populations from WT+WT vs WT+p53KO gastruloids depicted; *, p<0.05. **l**, Mean local density in mosaic gastruloids at 48h, 72h, or 96h as determined by the distance between 3D segmented viable nuclei. Analysis centered around clone-specific nuclei annotated in the figure legend. WT-mCherry (+WT), blue; WT-mCherry (+KO), green; WT-emiRFP (+WT), orange; p53KO-emiRFP (+WT), magenta. Mean of all nuclei from 3-5 gastruloids per condition. **, p<0.01. Corresponding to Extended Data Fig. 6g-h). **m-o**, Cytoskeletal and mechano-perception inhibition study. Absolute (**m-n**) and normalized (**o**) cell numbers for cell populations in WT+WT (**m**) or WT+p53KO (**n**) mosaic gastruloids at 120h, treated with 20 µM Rock inhibitor, 31 nM cytochalasin D, 5 µM Blebbistatin, or DMSO carrier control during 48-96h of gastruloid formation. n=18 gastruloids from 3 independent experiments. ****, p<0.0001; ns, not significant. All values depicted as mean ±SEM throughout this figure.

To directly test the influence of glucose availability on cell competition, we generated gastruloids in a pyruvate-poor N2B27 medium with varying glucose concentrations (**Fig. 4f-g; Extended Data Fig. 6c-d**). While the effect size of competition in this homemade medium was less pronounced than in commercial medium, lowering the amount of glucose eightfold from 20 mM to 2.5 mM did not alter cell competition (**Fig. 4h**). Counterintuitively, lowering the glucose levels even further to 0.5 mM, appeared to abolish the relative effect of cell competition. These data suggest that cell competition is not driven by glucose scarcity.

In *Drosophila*, zebrafish, and MDCK cells, cell competition can be mediated by reactive oxygen species (ROS) that selectively impair viability of the less fit loser cells ^31-34^. However, treating mosaic gastruloids with up to 500 μM of the ROS scavenger N-acetyl cysteine (NAC) did not alter cell competition, while higher concentrations impaired growth of all cell populations without changing relative cell competition effects (**Fig. 4i-k; Extended Data Fig. 6e-f**). This suggests that ROS does not mediate cell competition in mouse gastruloid.

We finally tested whether mechanical compression contributes to cell competition, as reported in *Drosophila* epithelial layers and mammalian 2D cultures ^17, 20^. Confocal microscopy and 3D segmentation of nuclei in mosaic WT+WT and WT+p53KO gastruloids detected lower local densities in gastruloids containing p53KO cells at 48h, no difference at 72h, and marginally higher densities in WT+p53KO gastruloids at 96h after aggregation (**Fig. 4l**). At 72h-96h the local density around early apoptotic WT-mCherry cells was elevated in WT+p53KO gastruloids compared to the average density around viable cells (**Extended Data Fig. 6g-h**). However, treatment with Rho-associated kinase inhibitors (ROCKi, Y-27632), cytochalasin D, or blebbistatin at highest sublethal concentrations did not significantly alter competition (**Fig. 4m-o**). Together, these results suggest that cell competition in 3D gastruloids is independent of size, glucose availability, ROS, or cytoskeletal tension, and is likely driven by other processes.

### Wnt and BMP signaling, but not ERK or Nodal signaling protect from cell competition

Relative differences or defects in BMP signaling have been reported to define loser cells ^5, 12^ and extrinsic elevation of BMP signaling has been shown to rescue loser cells in *Drosophila* dMyc supercompetition ^10^. Similarly, mispatterned cells that have too low or too high Wnt signaling for their relative position within developing zebrafish embryos get outcompeted by their neighbors via TGFb-SMAD signaling and cell-intrinsic ROS production and p53-independent apoptosis ^32^. Previous literature thus suggests a tight link between these signaling pathways and cell competition. We therefore set out to test whether modulating these and other signaling pathways in our system would influence the extend of cell competition (**Fig. 5a-c; Extended Data Fig. 7a-b**). Globally increasing BMP signaling by BMP4 treatment (1ng/mL at 48-72h) lowered the amount of cell competition when compared to BMP inhibition by LDN-193189 treatment (100nM at 48-72h) without affecting overall growth (**Fig. 5a-c**). ERK signaling activation or inhibition by FGF2 or PD0325901 (12.5 ng/mL or 1 μM at 48-72h), as well as modulation of ActivinA/Nodal/TGFb signaling by Activin A or SB431542 treatment (100 ng/mL or 10 μM at 48-72h) only altered cell competition slightly, but not significantly (**Fig. 5a-b**). Surprisingly, all tested signaling modulations resulted in more pronounced cell competition than observed in standard ChIR (CHIR99021, GSK3ß inhibitor; Wnt agonism) treated gastruloids (**Fig. 5c**), with DMSO control treatment allowing for the most drastic competition (**Fig. 5d-e**). The absence of ChIR treatment did however result in more poorly organized gastruloids that lacked Tbx6 expression and expressed elevated levels of Sox2, Otx2 and Sox3, indicating lack or delay of paraxial mesoderm development, and specifically in p53KO DMSO gastruloids a tendency toward more neuronal tissues (**Extended Data Fig. 7c-d**). Together, our observations suggested that Wnt agonism, similar to BMP but in contrast to ERK and Nodal signaling, might prevent cell competition, while absence of Wnt or BMP treatment is permissive for more extensive cell competition.

**Figure 5.**
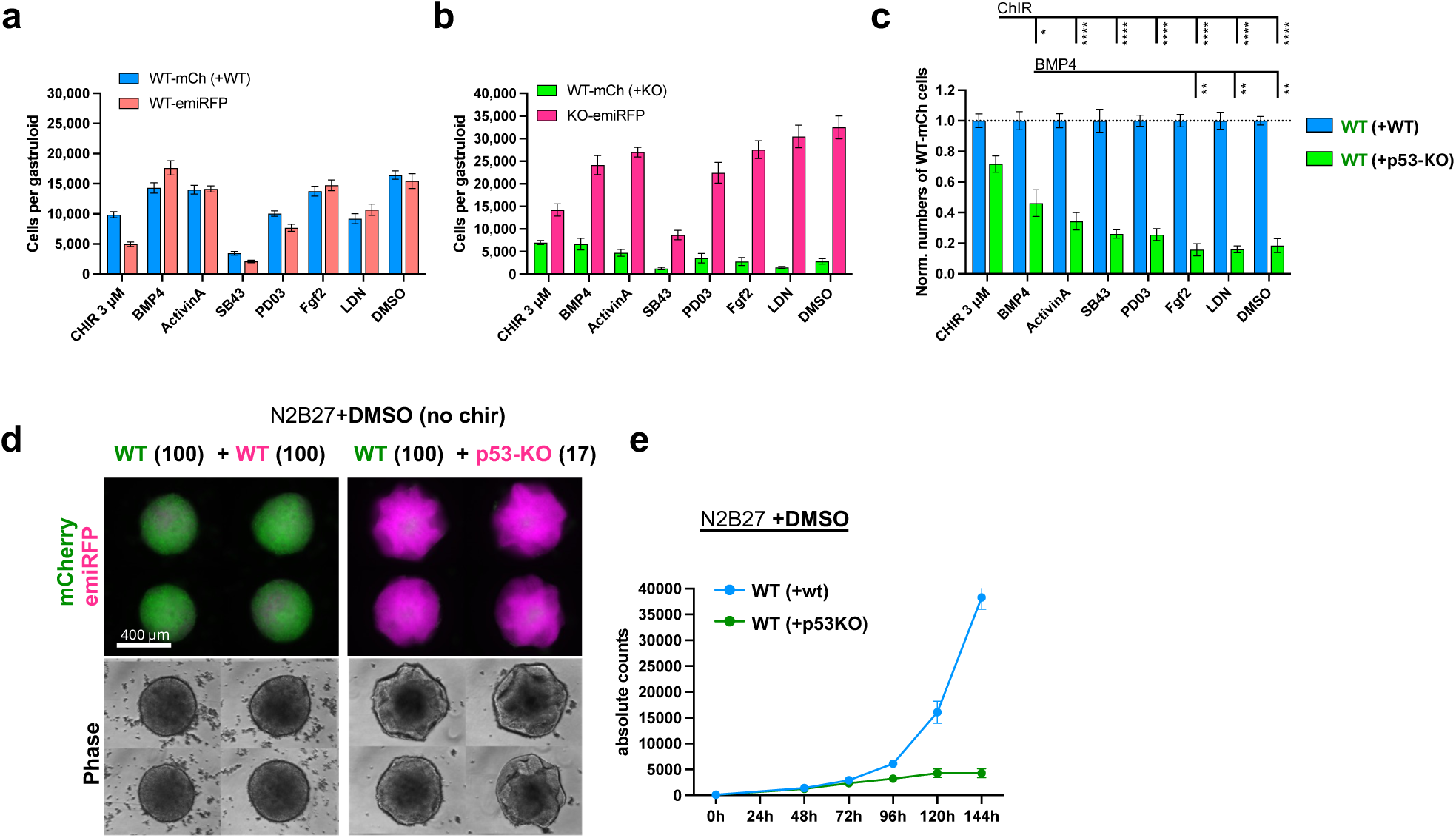
Wnt and BMP signaling, but not ERK or Nodal signaling protect from cell competition. **a-b**, Developmental signal perturbation during mosaic competition gastruloid growth. Absolute (**a-b**) and normalized (**c**) cell numbers for cell populations in WT+WT (**a**) or WT+p53KO (**b**) mosaic gastruloids at 120h. Gastruloids were treated with 3 µM ChIR (48-72h), 1ng/mL BMP4 (48-120h), 100ng/mL Activin A (48-72h), 10 µM SB431542 (48-120h), 1 µM PDO325901, 12.5 ng/mL FGF2 (48-120h), 100 nM LDN193189 (48-120h), or DMSO carrier control. n=12 gastruloids from 3 independent experiments. Only statistics between normalized WT-mCherry cell numbers in WT+p53KO gastruloids are depicted; *, p<0.05; **, p<0.01; ****, p<0.0001. **d**, Representative fluorescent micrographs and phase contrast images of 120h mosaic WT+WT (left) and WT+p53KO (right) gastruloids grown without ChIR treatment. Scale bar denotes 400 µm. **e**, Absolute growth curves of WT-mCherry cells in mosaic gastruloids without ChIR treatment (DMSO control, as depicted in panel **d**) seeded together with WT-emiRFP (blue) or p53KO-emiRFP (green) cells. n=9 gastruloids per datapoint from 3 independent experiments. All values depicted as mean ±SEM throughout this figure.

### Cell competition is differentiation stage specific

At 48h after aggregation, gastruloids have exited naïve pluripotency and entered a formative-like state (**Fig. 6a-b**). The standard gastruloid protocol calls for a ChIR treatment (Wnt agonism) at 48-72h after aggregation (**Fig. 1c**), which coincides with the onset of cell competition and the entry into gastrulation. Recent work has demonstrated that this Wnt signaling pulse at 48h not only drives development of the primitive streak but also temporally accelerates its developmental progression ^35^. Indeed, mosaic gastruloids dissociated, stained for the pluripotency surface markers SSEA-1 and Pecam1, and analyzed by flowcytometry support that the exit of pluripotency is delayed in the absence of ChIR (**Fig. 6c-d; Extended Data Fig. 8a-b**). DMSO control treated gastruloids display the same SSEA-1 intensity at 96h as at 48h (**Fig. 6c, upper panel**) while ChIR treated gastruloids almost entirely lose SSEA-1 positivity at 96h (**Fig. 6c-d; Extended Data Fig. 8a-b**). These data propose that the absence of ChIR treatment delays gastruloid differentiation and might extend the time that gastruloids remain in the developmental stage permissive for cell competition, which could cause the observed increase in cell competition magnitude (**Fig. 5d-e**).

**Figure 6.**
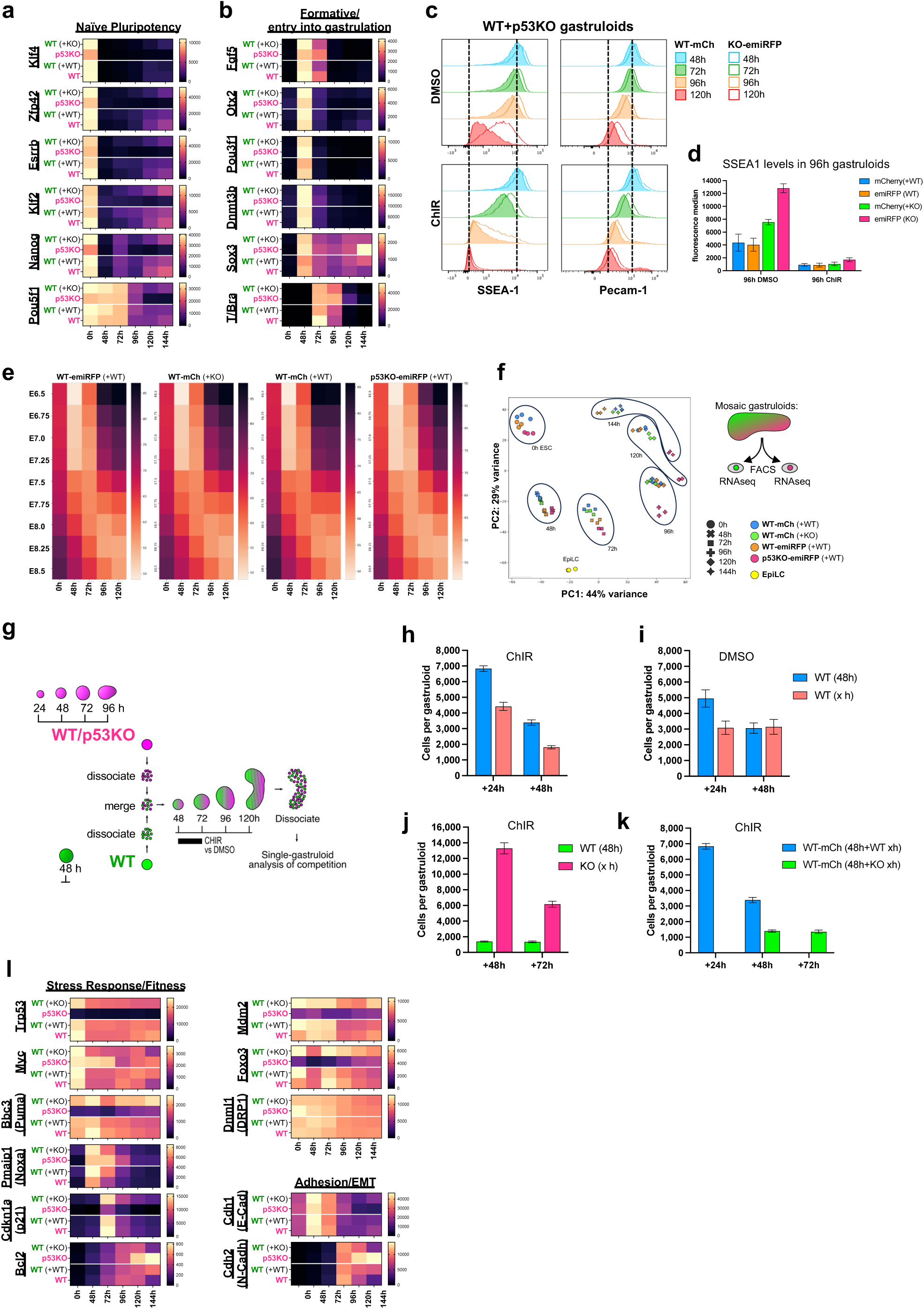
Cell competition is differentiation stage specific, but does not depend on relative shift in pluripotency. **a-b**, Heatmap of temporal gene expression dynamics of markers of naïve pluripotency (**a**), formative pluripotency (**b**), or entry into gastrulation (**b**). FACS-isolated cell populations from mosaic gastruloids depicted separately in rows and timepoints from 48h-144h after aggregation in columns. Each datapoint is a mean of 3 biological replicates. **c**, Flow cytometric quantification of pluripotency surface marker expression SSEA1 and Pecam-1 in WT-mCherry (filled histograms) and p53KO-emiRFP (open lines) from mosaic gastruloids treated with ChIR (lower panels) or DMSO control (upper panels) analyzed at 48h, 72h, 96h, or 120h. **d**, Quantification of SSEA-1 surface marker expression in mosaic gastruloids at 96h as determined by flow cytometry depicted in panel c. n=3 independent experiments. **e**, Temporal alignment of gastruloid and mouse embryo development. Heatmap of Euclidean distance between transcriptomes of FACS-isolated populations from mosaic gastruloids and pseudobulk transcriptomes of mouse embryos at indicated developmental timepoints. Each datapoint is the mean of 3 biological replicates. **f**, Principal component analysis comparing FACS-isolated cell populations from mosaic gastruloids from 48h, 72h, 96h, 120h, 144h after aggregation, together with 0h mESC clones pre-gastruloid aggregation, and Epi-like cells (EpiLC). **g**, Schematic of heterochronic mosaic gastruloid formation protocol. **h-i**, Heterochronic mosaic gastruloids created from 48h WT-mCherry gastruloid cells and 24h or 48h WT-emiRFP gastruloid cells in standard ChIR conditions (**h**) or DMSO control (**i**). Bar graphs depict absolute cell counts of populations in 120h gastruloids, 72h after reaggregation. n=12 gastruloids from 3 independent experiments. **j**, Heterochronic mosaic gastruloids created from 48h WT-mCherry gastruloid cells and 48h or 72h p53KO-emiRFP gastruloid cells in standard ChIR conditions. Bar graphs depict absolute cell counts of populations in 120h gastruloids, 72h after reaggregation. n=12 gastruloids from 3 independent experiments. **k**, Comparative graph of WT-mCherry absolute counts from varying heterochronic conditions detailed in panels **h-j**. n=12 gastruloids from 3 independent experiments. **l**, Heatmap of temporal gene expression dynamics of markers of stress response and apoptosis, cellular fitness, and EMT/adhesion. FACS-isolated cell populations from mosaic gastruloids depicted separately in rows and timepoints from 48h-144h after aggregation in columns. Each datapoint is a mean of 3 biological replicates. All values depicted as mean ±SEM throughout this figure.

The flowcytometry analysis of SSEA-1 and Pecam1 levels additionally revealed that within each timepoint, p53KO cells expressed higher levels of pluripotency markers than their WT counterparts and lagged behind in their downregulation (**Fig. 6c, open vs filled histograms, Extended Data Fig. 8b**), while non-autonomously delaying neighboring WT-mCherry cells in mosaic gastruloids (**Extended Data Fig. 8b**). A similar developmental delay was observed in the transcriptional downregulation of Brachyury after 72h and Myc after 48h despite the presence of ChIR (**Fig. 6b,l**). However, on a whole transcriptome level, all four populations from mosaic WT+WT or WT+p53KO gastruloids followed a similar temporal trajectory as measured by Euclidean distance to times embryonic transcriptomes (**Fig. 6e**). Principal component analysis of sorted clones from mosaic gastruloids indicated that p53KO cells cluster closely to the other clones until 72h, after which they diverge and separate from the remaining 3 cell populations (**Fig. 6f**). Together, these data suggest that the developmental delay observed in p53KO cells might be specific to pluripotency networks, rather than overall developmental timing.

We thus set out to interrogate whether this developmental delay of the supercompetitor cells was necessary and/or sufficient for cell competition by creating heterochronic mosaic gastruloids. To do this, gastruloids of the individual clonal lines were generated, dissociated, and re-aggregated at different timepoints, centered around the 48h timepoint of WT-mCherry cells in question (**Fig. 6g**). Introducing WT-emiRFP cells from 24h gastruloids into WT-mCherry 48h gastruloids did not impair their growth, but supported it, indicating that a relative difference in pluripotency/developmental stage between two WT clones is not sufficient to cause cell competition (**Fig. 6h**). The same was observed in heterochronic WT+WT DMSO gastruloids that potentially allowed for a longer sustained heterochrony of WT cells (**Fig. 6i**). Reintroducing p53KO cells from 72h gastruloids into 48h WT gastruloids caused almost identical cell competition pressure on the WT cells as re-introducing homochronic 48h p53KO cells (**Fig. 6j-k**). This suggests that a 24h relative difference in developmental progression is neither necessary not sufficient for cell competition. However, conducting an expanded panel of heterochronic mosaic gastruloids revealed that 96h p53KO cells no longer exert any competition pressure on neighboring 48h WT-mCherry cells, indicating that supercompetition requires both cell populations to reside within the developmental window of time permissive for cell competition, which is centered around 72h (**Extended Data Fig. 8c-e**). Taken together, p53KO cells display a relative developmental delay during gastruloid formation, but this developmental delay is neither necessary nor sufficient to cause cell competition. Instead, cell competition relies on both competing cells to reside within a developmental window of time permissive of cell competition, which can be shortened or prolonged by Wnt agonism in gastruloids.

We next sought to understand the temporal regulation of cell competition from a transcriptional point of view. Previous work by others suggested downregulation of DRP1 (*Dnml1*) ^36^, and p53-dependent upregulation of pro-apoptotic Puma (*Bbc3*) expression during primed pluripotency ^14^ as causes for hypersensitivity to apoptosis. We too observed in mosaic gastruloids that *Dnml1* is downregulated during differentiation and that winner cells express higher levels than loser cells within the same gastruloid (**Fig. 6l**). While we confirm that Puma is expressed highest at 48h of gastruloid formation, we observe that pro-apoptotic Noxa (*Pmaip1*) is upregulated at 48h with even sharper temporal specificity (**Fig. 6l**). Unexpectedly, p53KO cells also upregulate Noxa sharply at 48h of gastruloid formation and maintain its expression longer than WT cells (**Fig. 6l**), hinting toward a p53-independent transcriptional regulation of Noxa. This upregulation of Puma and Noxa is followed by upregulation of the p53-dependent stress response gene p21 (*Cdkn1a*) 72h after aggregation, which marks the temporal center of cell competition. Anti-apoptotic Bcl2 is upregulated from 96h onward (**Fig. 6l**), coinciding with gastrulation and a switch from E-Cadherin to N-Cadherin (**Fig. 6l**). Together, these temporal expression dynamics suggest that the stress response machinery is highly expressed during primed pluripotency, but that p53-p21 dependent stress is only activated 24 hours later at the onset of gastrulation, and that apoptosis is temporally limited by the downregulation of the apoptosis machinery and upregulation of Bcl2.

### EpiGastruloids do not undergo cell competition

Our experiment of preventing ESC aggregates from exiting pluripotency confirmed that cell competition does not take place in absence of differentiation, even in a 3D context, but instead only after passing through formative pluripotency. This led us to test whether 3D cell aggregates still undergo cell competition if aggregated at a later developmental stage, purposely missing our proposed developmental window of cell competition. Toward this, we generated mosaic competition Epi-gastruloids ^37^. mESCs were first converted into an epiblast-like state during 3 passages in N2B27+AA+bFGF on fibronectin treated wells. These epiblast-like cells (EpiLC) are transcriptionally comparable to 48-72h gastruloid aggregates (**Fig.6f**) and are likely in a formative state of pluripotency (*high expression of Fgf5, Otx2, Oct6, Sox3, Dnmt3b*). Prior to 3D aggregation, the cells were cultured 48h without Activin A and 24h with ChIR, comparable to the ChIR pre-treatment of human gastruloids ^38^. At the time of 3D aggregation, these cells transcriptionally resemble the mouse caudal epiblast ^39^ and are thus referred to as Epi-CE cells (**Fig. 7a-b**). 72h after aggregation, these Epi-gastruloids reach a developmental stage equivalent to 120h classical gastruloids (**Fig. 7c-d**). However, compared to classical gastruloids at 120h, 72h Epi-gastruloids show an enrichment in *Neurog2*, *Sox1*, *Sox2*, *Foxg1*, *En1* which might indicate preferential spinal cord and brain trajectories (**Fig. 7c-d**), while many other lineages seem underrepresented. Immunofluorescent staining of 72h Epi-gastruloids confirmed this by absence of endoderm and mesoderm markers, but abundant expression of Sox2 and Sox3 (**Extended Data Fig. 9a-b**). Hox gene expression in Epi-gastruloids is shifted toward posterior late hox genes (less *Hoxa1*, *Hoxb1*, *Hoxd1* expression, more *Hoxa10*, *Hoxc10*, *Hoxa11* expression) when compared to classical 120h gastruloids (**Fig. 7e-f**). Together these data suggest that Epi-gastruloids might be a 3D developmental system modeling late gastrulation starting from caudal epiblast and leading to posterior neuroepithelial like development.

**Figure 7.**
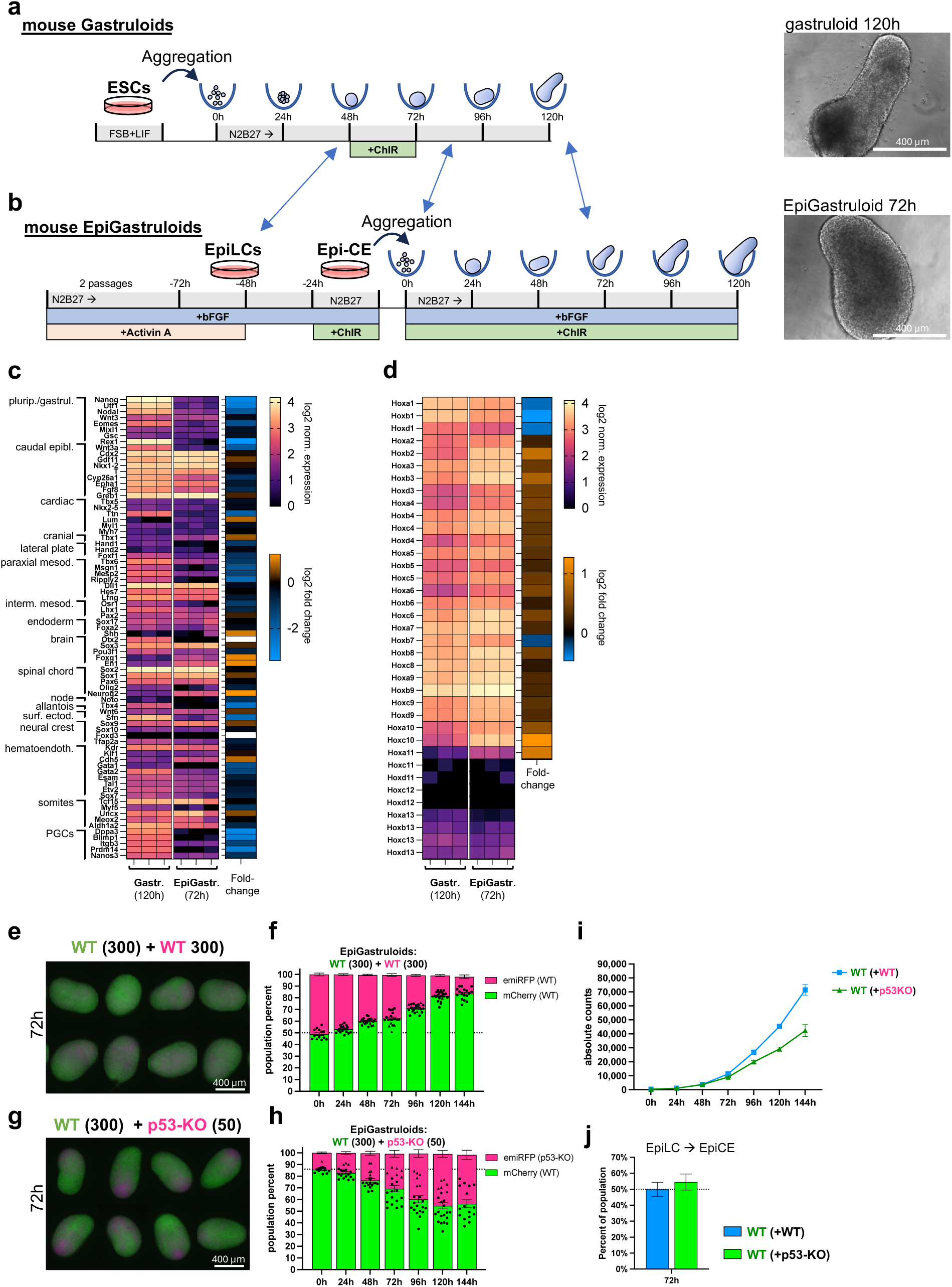
EpiGastruloids compete less than gastruloids. **a-b**, Schematic experimental protocol of mouse gastruloid (**a**) and mouse EpiGastruloid (**b**) formation and representative example brightfield images of 120h mouse gastruloid (top) and 72h EpiGastruloid (bottom). EpiGastruloids are generated by transitioning mESCs into Epi-like cells (EpiLC), followed by a treatment resulting in caudal epiblast-like cells (EpiCE) and lastly, 3D aggregation. Blue arrow point out tentative developmental stage equivalence between standard mouse gastruloids and mouse EpiGastruloids. **c**, Heatmap of developmental lineage marker expression profiles from mRNA bulk sequencing analysis of 120h WT mouse gastruloids and 72h WT mouse EpiGastruloids. **d**, Heatmap of Hox gene expression of 120h gastruloids and 72h EpiGastruloids. In panels **c** and **d,** log2 normalized transcript levels are depicted as magma heatmap and log2 fold-change of EpiGastruloids compared to standard gastruloids are depicted as adjacent blue-to-orange heatmap. **e-h**, Fluorescent widefield micrographs of WT+WT (**e**) and WT+p53KO (**g**) EpiGastruloids at 72h and corresponding single EpiGastruloid flow cytometry analysis (**f, h**) depicting population composition as percentage of mCherry+ (green) and emiRFP+ (magenta) cells over time. n=13-20 EpiGastruloids from 4 independent experiments. **i**, Absolute growth curves of WT-mCherry cells in mosaic EpiGastruloids seeded together with 300 WT-emiRFP (blue), or 50 p53KO-emiRFP (green) cells. n=17 gastruloids per datapoint from 3 independent experiments. **j**, Percentage of WT-mCherry cells after co-transitioning from EpiLC to EpiCE together with WT-emiRPF (blue) or p53KO-emiRFP (green) cells. n=3 independent experiments. All values depicted as mean ±SEM throughout this figure. All scale bars denote 400 µm.

In contrast to our observations in classical gastruloids, mosaic Epi-gastruloids do not undergo the same pronounced cell competition, but instead only display a slow drift between cell populations (**Fig. 7e-i**). The co-cultures during the 2D transition stage from EpiLC to EpiCE do not undergo competition (**Fig. 7j**). Together, these data indicate that Epi-gastruloids model later and more posterior stages of development compared to classical gastruloids, and that cell competition is restricted to gastruloids undergoing the transition of formative epiblast to gastrulation.

## Discussion

Early embryonic development is a highly orchestrated process in which cells build and develop a body plan through a spatiotemporal coordination of mechano-chemical interactions within multicellular ensembles. A key event in this is gastrulation, the process whereby the mass of cells that results from the proliferation of the zygote, outlines the body plan ^1^. However, the events leading up to gastrulation are not without complications; 73% of human pre-implantation embryos are mosaics containing aneuploid cells ^40^ and 8-33% of conceptions are lost within the first three weeks ^41-43^. Furthermore, in all mammals, gastrulation is associated with a dramatic increase in cell proliferation that requires high cellular fitness. Embryos must therefore employ rigorous and timely mechanisms of quality control that ensure only the fittest cells within the epiblast contribute to the final body plan, while damaged or suboptimal cells are removed ^6-8^. Here, we use gastruloids as an easy to perturb 3D *in vitro* model of gastrulation to study the process of cell competition.

We observe that in gastruloids, p53KO cells act as supercompetitors and cause apoptosis of neighboring WT cells, while WT+WT mosaics coexist without competition. This is in agreement with previous observations in adherent culture ^18^ but here this is extrapolated to an embryo-like 3D situation. Our results show that different clonal WT populations of equal competitive fitness do not exert any influence on each other’s growth behavior, regardless of their ability or inability to undergo apoptosis, and irrespective of their absolute number of neighbor cells (**Fig. 1h-j**; **Fig. 3f-j**). In contrast, seeding as few as two p53KO cells per mosaic gastruloid (1.3% of the starting cell population) is sufficient to measurably impair growth of neighboring WT cells (**Fig. 1f-g**), and 25 p53KO cells robustly outcompete 150 WT cells without exception. These observations demonstrate an extensive effect size of cell competition in our 3D gastruloid system and validate it as a highly suitable model to study this phenomenon.

The ability of p53KO cells to outcompete WT cells at such disproportionate ratios raises the intriguing question of whether, in embryos, emergence of single p53-mutant clones before gastrulation invariably results in monoclonal p53-mutant embryos post-gastrulation.

Blc2 is a dominant negative inhibitor of apoptosis and overexpression confers apoptosis resistance. However, we report that even large numbers of Bcl2-overexpressing WT cells exert no cell competition pressure on neighboring WT cells, highlighting that the winner cell status of p53KO supercompetitor cells is not simply a result of their resistance to apoptosis. We believe that lack of the stress sensor p53 causes cells to perceive their own fitness as unrestricted, thereby eliciting their winner status in relative fitness comparisons. Loser cells respond to fitter cells by stabilizing p53 protein, which results in their apoptosis ^5, 17^. This intriguingly positions p53, or lack thereof, as a fitness-defining factor upstream of cell competition and simultaneously downstream of the fitness comparison as part of the loser cell response.

The interactions between WT and p53KO supercompetitor cells go beyond a one-sided growth impairment. While compensatory growth in response to apoptosis has been demonstrated in *Drosophila* ^44^, and suggested in mouse embryos, where cell competition does not influence the final embryonic size ^11^, recent supercompetition studies in 2D stem cell differentiation models have proposed that winner cells do not respond to apoptosis of loser cells ^18^. However, mosaic gastruloids reveal clear compensatory growth of p53KO cells that depends on the apoptosis of neighboring WT cells (**Fig. 1k; Extended Data Fig. 4g**), suggesting that cell competition entails two-way communication between competing cells. These observations underscore potential differences between 3D and 2D systems, and suggest that, while fitness is measured in diverse contexts, the respective mechanism of competition might be context dependent.

Flow cytometry revealed extensive phagocytic uptake of debris from apoptotic neighbors, suggesting that nutrient recycling in winner cells may support this compensatory growth. Additionally, we observed that p53KO cells exerted cell non-autonomous effects on neighboring WT cells’ development. p53KO cells in gastruloids were enriched for paraxial mesoderm, intermediate mesoderm and caudal epiblast fates, and conferred that same trend onto their neighboring WT cells in mosaic gastruloids, albeit to a lesser extent (**Extended Data Fig. 2e; Fig. 6f**). However, despite this enrichment trend, the transcriptomes of WT(+WT) and WT(+p53KO) cells were still highly similar and indicate that no essential lineages were lost during cell competition. This suggests that despite the significant effect size of WT cell elimination, cell competition in gastruloids does not target specific lineages preferentially and is largely phenotypically silent in terms of cell fate specification.

Previous studies proposed that relative pluripotency levels, as a readout of developmental timing, might be a determinant of winner or loser status during cell competition ^13, 19^. Myc expression levels, which decrease over developmental time, could be the functional link to homogenize pluripotency levels and timing across the embryo ^13^. Our transcriptome data from mosaic gastruloids confirm that Myc expression is progressively downregulated during differentiation, but not in p53KO cells which maintain prolonged high Myc expression (**Fig. 6c**). In accordance, we find that p53KO cells also downregulate other pluripotency markers with a delay compared to WT cells, fitting to above mentioned hypothesis. However, the modular nature of gastruloids uniquely enabled us to functionally test the relation of pluripotency and competitive fitness by creating heterochronic mosaic gastruloids, which contain populations of differing genotypes and developmental stages. Unexpectedly, we find that relative shifts in pluripotency are neither necessary for p53KO supercompetition, nor sufficient to convey a competitive advantage among different WT populations (**Fig. 6g-k**). Together this highlights that a relative difference in pluripotency levels between WT and p53KO cells exists, but that this may not play as much of a role, as the presence of both competing cell populations within the developmental window permissive for cell competition.

We tested the influence of morphogen signalling in cell competition in gastruloids and observe a clear effect of Wnt signalling in this process. It has been suggested that Wnt signaling accelerates the progression from formative pluripotency to gastrulation corresponding to gastruloid at 48h to 96h and mouse embryos at E5.5 to E7.5 ^35^. Here, we show that in the absence of a ChIR pulse, gastruloids reside longer in the developmental window permissive for cell competition, which results in a more drastic competition effect size (**Fig. 5d-e**). This indicates that the temporal window of cell competition correlates to a developmental stage, which can be artificially prolonged or shortened *in vitro*. The temporal restriction of cell competition to this window of development may partially be due to a downregulation of DRP1 (*Dnml1*) during primed pluripotency, which causes hypersensitivity to apoptosis ^36^, and to the temporal upregulation of the stress response and pro-apoptotic machinery. Our time-resolved transcriptomic analysis in mosaic gastruloids draw a timeline in which cells prepare an apoptotic response machinery through expression of pro-apoptotic genes during primed pluripotency, but only experience p21-mediated stress response 24h later during the onset of gastrulation, after which stress response genes are downregulated and pro-survival *Bcl2* expression increases (**Fig. 6l**). This may be a mechanism to temporally limit cell competition to pre-gastrulation stages, where lost cells can be replaced without compromising germ layer and cell lineage specification. Previous studies had already suggested the temporal p53-dependent upregulation of *Puma* expression during primed pluripotency ^14^, which we confirm, but we additionally observe a temporally more specific upregulation of *Noxa*, which unexpectedly is p53 independent.

Apoptosis in the mouse embryo peaks just before gastrulation, after which it is markedly reduced. Similarly, it is often assumed that cell competition ends at gastrulation, but without it having been appropriately tested. It had not been clear whether cells are unable to compete after gastrulation, or whether early cell competition simply homogenized the epiblast sufficiently to prevent later cell competition. To test directly whether cells lower their sensitivity to cell competition after entry into gastrulation, we generated mosaic EpiGastruloids (modified after: ^37^), in which cells of varying fitness levels only encounter each other after having reached a developmental equivalent of 72-96h gastruloids (**Fig. 7**). For this, we transitioned ESCs into Epi-like cells (EpiLC, formative pluripotency), a transition state towards Epi-stem cells (EpiSCs, ∼E7.5; ^45^). The EpiLC were further transitioned into caudal epiblast-like cells (Epi-CE, late gastrulation), followed by mosaic aggregation of varying clones into 3D EpiGastruloids. Unlike gastruloids, EpiGastruloids show almost no cell competition within the first 72h, suggesting that cells are unable to compete after the onset of gastrulation. The changed growth behavior of WT-mCherry cells after 72h of EpiGastruloid formation could be attributed to the formation of neuronal lineages (brain and spine), which might undergo a second and separate wave of cell competition ^46^, but this remains to be investigated in detail in the future.

In addition to Wnt signalling, we find that BMP signaling also reduces cell competition at the onset of gastrulation. This draws intriguing parallels to a recent study showing that increased Wnt and BMP signaling lower the amount of chromosome missegregation and DNA replication stress in pluripotent stem cells, while inhibition of Wnt and BMP, or activation of FGF signaling, increases it ^47^. In our system activation of FGF signaling has slight effects of increasing cell competition. Together, this suggest that these signaling pathways may modulate DNA replication stress and cell competition in a coordinated manner, thereby focusing quality control toward cell populations that require it most. This correlation of cell competition to anteriorizing and posteriorizing signals, raises the question whether cell competition displays local differences along the antero-posterior axis in the epiblast. Signaling dependent effects on chromosome missegregation, as well as cell competition, diminish at gastrulation and both reemerge during early brain development ^46, 47^ hinting toward a broader link between chromosome segregation fidelity and cell competition sensitivity. Alternatively, global Wnt or BMP agonism could also lower cell competition by overwriting other developmental programs, thus homogenizing the cells within gastruloids and lowering heterogeneity.

Extrinsic apoptosis is triggered by external signaling through FasL/Fas interactions in the TNF receptor family. Through Fadd adaptor proteins caspase 8 gets activated and either directly result in apoptosis, or via Bid-cleavage cross activates intrinsic apoptosis. Bcl2 expression rescues intrinsic apoptosis and partially mitigates extrinsic apoptosis. Previous studies have demonstrated that cell competition can be prevented by Bcl2 overexpression, but also that caspase 8 is strongly upregulated as a function of cell competition and that cell competition might be mediated via short range or direct contact of cells ^18^, which left the question whether cell competition is mediated by intrinsic or extrinsic apoptosis. We confirm that Bcl2 overexpression fully prevents cell competition, that TNF ligands are not expressed within the competition timeframe, and that knockout of *Fas*, *FasL*, or *Fadd* does not affect cell competition, adding to our confidence that cell competition in gastruloids is exclusively mediated by intrinsic apoptosis. This finding distinguishes developmental cell competition from other systems like the hair follicle stem cell niche, where competition relies on extrinsic apoptosis signaling ^26^.

In summary, we establish 3D gastruloids as a robust platform to study cell competition, overcoming limitations of 2D models while enabling precise spatiotemporal analysis. By leveraging the modular and perturbable nature of this system, we provide key insights into the temporal and mechanistic regulation of cell competition, and create a foundation for future investigation into embryonic quality control and its implications for developmental biology and disease.

## Acknowledgments

We thank Tristan Rodriguez for sharing a template for the Bcl2 overexpression plasmid used in this study. Andre Dias has helped with discussion and advice on developmental marker genes. Vikas Trivedi and the EMBL Barcelona mesoscopic imaging facility have provided access to imaging equipment vital to this study. The gRNA used to create a *Trp53*-knockout was a gift from Ana Janic, who has also generously given us access to the materials and apparatus for Western blotting. We thank the UPF and CRG Flow Cytometry, Advanced Light Microscopy, and Genomics core facilities for support and instrumentation for this project. We thank Maria Constantina Balayo Costal who has been a constant support throughout the project.

## Methods

### Cell culture

E14Tg2A mouse embryonic stem cells (ESCs) were cultured at 37C in 5% CO2 in ES/LIF medium on gelatin coated dishes unless otherwise stated. ES/LIF medium was composed of GMEM (Thermo Fisher Cat# 11710-035) supplemented with 15% Fetal Bovine Serum (FBS; Thermo Fisher Cat#10270106), 1xNon-Essential Amino Acids (Thermo Fisher Cat# 11140-035), 1mM Sodium Pyruvate (Thermo Fisher Cat# 11360-039), 2mM L-Glutamine (Thermo Fisher Cat# 25030-024), 0.1 mM 2-mercaptoethanol (Thermo Fisher Cat# 31350010), 10ng/mL LIF (Qkine Cat# Qk018-0100). ESCs were passaged as single cells every 2 days and medium was changed every day. Cells were routinely tested negative for mycoplasma infections.

### Generation of clonal cell lines

All stable cell lines were generated from single clones using the piggybac transposase system ^48^ and E14Tg2A cells. mESCs at passage 12-15 were co-transfected with pBase piggybac transposase expression plasmid and the piggybac donor plasmid of interest using Lipofectoamine 3000 following the manufacturers protocol. One week after transfection, when transient expression of the transfected plasmids had resided, cells with stably inserted fluorophore expression were sorted by fluorescence activated cell sorting (FACS) into 96 well plates as one cell per well. Clonal colonies were expanded and tested for the ability to form gastruloids. The H2B-emiRFP670 p53 knock-out clone “A12-8” is a direct daughter clone of the H2B-emiRFP670 WT clone “A12”. For this, A12-WT cells were co-transfected with a FUCas9Cherry expression plasmid (Addgene #70182) and a plasmid carrying an sgRNA targeting TP53 *5′-GGCAACTATGGCTTCCACCT-3′*, as previously described ^49^. Transfected cells were selected for successful p53 loss-of-function by treatment with 10 μM Nutlin-3a for 7 days. Single cell clones were generated from the Nutlin3-a resistant cells, which were subsequently tested for p53 knock-out by Western blot and genomic locus sequencing. The clone A12-8, which was selected for this study is Nutlin-3a resistant, has no p53 protein as detected by western blot, and carries a homozygous +1 frameshift insertion in exon 4 of the TP53 gene.

### Western blot confirmation of p53-KO clones

P53 knockout candidate clones were treated with 5 μm Nutlin-3a for 24 hours to prevent MDM2-mediated p53 degradation. Whole cell lysates were generated and analyzed by Western blot as previously described ^50^ under omission of the DDM detergent. In short, cells were lysed in RIPA buffer supplemented with complete protease inhibitors (Roche, Cat#11836170001). Chromatin was sheared by sonication and lysates were precleared by centrifugation at 15,000 xg for 10 minutes at 4°C. Protein content was determined using a Protein assay kit (BioRad, Cat# 500-0006). Protein samples were boiled and reduced in Laemmli buffer with β-mercaptoethanol, and 5 μg protein per sample were separated by SDS-PAGE. After transfer onto nitrocellulose membrane, membranes were analyzed by ponceau staining, and blocked with 5% milk powder in PBS-T (PBS containing 0.1% Tween 20). Primary antibodies against p53 (1:1,000; COMPANY, Cat# XXX) and β-actin (1:2,000; Santa Cruz, Cat# sc-47778) were incubated overnight at 4°C. HRP-conjugated secondary antibodies were incubated for 1 hour at room temperature, and bands were visualized using the ECL Prime chemiluminescence system and imaged on a ChemiDoc MP (BioRad).

### Mosaic Gastruloid formation

Gastruloids were generated as previously described (Beccari et al., 2018) unless otherwise mentioned in the manuscript. ESCs grown in ES/LIF medium (described above) were seeded as 45,000 cells per 6-well dish 3 days prior to gastruloid formation and medium was exchanged daily. On the day of gastruloid seeding, cells were washed with PBS -Ca^2+^/-Mg^2+^ (PBS-/-) and enzymatically dissociated using prewarmed trypsin for 1.5 minutes. Cells were gently pipetted up and down to form a single cell suspension, and Trypsin was washed out using an excess of ES medium followed by pelleting the cells at 200xg for 2 minutes. Cells were washed twice more with PBS-/- and finally resuspended in N227 Diff (NDiff227-custom, Takara, Cat# XA0530). Cells were counted and resuspended in a cell master mix that would yield the desired number of cells per gastruloid in 40 µL per U-bottom 96-well (Ultra-low attachment plates Grainer Cat#650970). For single culture gastruloids 300 cells were seeded per gastruloid, with exception of p53-KO cells, for which seeding 50 cells yield a gastruloid of comparable size at 120 hours after aggregation. For mosaic gastruloids, 50% of these numbers were used, resulting in 150+150 cells of WT:WT cell mosaics, and 150+25 cells of WT+KO mosaic gastruloids. Gastruloids were not perturbed for the first 48h after aggregation. At 48h, 150 µL of fresh N2B27 containing 3µM of ChIR (Chi99021, Sigma Cat#SML1046-5MG) were added to the gastruloids. At 72h, 96h, and 120h each, 150 µL of medium was removed from the wells, and fresh 150 µL N227Diff were added.

### Microscopy

Epifluorescence widefield microscopy of the H2B-tagged fluorophores in gastruloids and 2D cultures was conducted on live cells at 37°C and 5% CO_2_ using a Zeiss AxioObserver.

Confocal imaging and immunofluorescence staining were conducted on fixed gastruloids. Gastruloids were harvested at indicated time points and washed once in PBS +Ca^2+^/+Mg^2+^ (PBS+/+) followed by fixation in 4% paraformaldehyde (PFA, Aname EMS, Cat#15710) overnight at 4°C. Gastruloids we washed 3 times in PBS + 0.2% Triton X-100 (Sigma Cat#T8787)(PBS-T) the following day and blocked in PBS-T + 10% Bovine Serum Albumin (BSA, Merck Life Science, Cat#3117332001) for 1.5 hours at room temperature. Primary antibodies were diluted in PBS-T + 2% BSA and incubated with gastruloids at 4°C overnight under agitation. Gastruloids were washed 6 x 10 minutes in PBS-T the following day. Fluorophore-conjugated secondary antibodies were diluted 1:500 in PBS-T + 2% BSA together with 1 μg/mL DAPI (Thermo Fisher Cat#D1306) and incubated with gastruloids at 4°C overnight under agitation. Gastruloids were washed 6 x 10 minutes in PBS-T the following day and mounted on microscopy slides. For mounting, thin strips of double-sided tape were placed onto the microscopy slides forming a square to create spacers. Gastruloids were placed onto the microscopy slide between the spacers, PBS was removed and gastruloids were covered in a drop of Vectashield antifade mounting medium (Palex Vector Cat# H-1000-10) a thin coverglass was placed ontop of the gastruloids and the spacers without excerting force onto the gastruloids. Microscopy was conducted using a Zeiss LSM980 confocal laser scanning microscope.

### Flow cytometry from gastruloids

Gastruloids were collected at indicated timepoints, washed in PBS -/-, and enzymatically dissociated using Accutase (Lab Clinics Capricorn Scientific, Cat#ACC-1B). Optimal dissociation was achieved by warming tubes containing gastruloids and Accutase to 37°C in a water bath while shaking and eventually flicking the tubes during a time course of 5 minutes. Gastruloids were further dissociated mechanically using a p1000 pipette while adding flow buffer (PBS-/-, 2% BSA, 2mM EDTA). Resulting single cell suspensions were washed in flow buffer and pelleted by centrifugation at 200 xg for 3 minutes. Single gastruloid flow as depicted in Figure 2 of this manuscript was conducted by analyzing the dissociated gastruloids live at this step of the protocol. For antibody stained flowcytometry multiple gastruloids of the same time point and batch were pooled.

For antibody staining, single cells were washed one more time in PBS-/- followed by fixation in 4% PFA for 15 minutes. Fixed cells were washed 3 times in flow buffer, followed by blocking in 10% BSA in PBS-/- containing 0.1% Triton X-100 for 30 minutes at room temperature. After setting aside a fraction of the sample for use as negative or secondary antibody control, primary antibodies were diluted in 2% BSA in PBS-/- with 0.1%Triton X-100 (staining buffer). Cells were incubated with primary antibodies for 1 hour shaking at room temperature, followed by 3 washing steps in staining buffer. Fluorophore-conjugated secondary antibodies were diluted 1:500 in staining buffer unless otherwise stated and used to stain cells for 30 minutes at room temperature shaking. Cells were washed 3 times in staining buffer again and transferred to flow cytometry tubes for analysis. All samples were analyzed using a BD Bioscience LSRFortessa system.

### Absolute counts of cells within gastruloids

The method for generating absolute counts of cells per gastruloid was adapted after (Merle…Gregor 2024). Individual gastruloids were transferred to Eppendorf tubes and washed in PBS -/-. After aspirating the supernatant, Accutase was briefly prewarmed at 37°C for 45 seconds and directly added to each gastruloid at 50 µL per tube. After 5 minutes incubation, the 50 µL of Accutase containing the enzymatically loosened but not yet disintegrated gastruloid was transferred to a 96-well flat bottom PhenoPlate (Perkin Elmer, Cat#6055302) and pipetted up and down 20 times. Gastruloids of early time points disintegrate very easily into single cells, but later time points 120h or 144h gastruloids might require additional pipetting after visual inspection. Pipette tips for this step should be pre-wetted in Accutase to prevent losing single cells to stickiness in the tip. The single cells were then fixed by adding 100 µL of 1% PFA in PBS -/- per flat bottom well. Gastruloids of different time points were sequentially dissociated and fixed to generate growth curves of individual endpoints. The PhenoPlates were then tile-imaged using a Perkin Elmer Opera Pheonix to detect the nuclear H2B-mCherry or H2B-emiRFP670 of the varying cell populations. Tiled images were stitched, and nuclei were automatically segmented counted in high throughput using a custom ImageJ macro. For each condition or time point, a minimum of 4 gastruloids were used as technical repeats, and all conditions were repeated in multiple independent experiments as annotated.

### 3D image segmentation

3D cell segmentation was performed using the QLIVECELL software (https://github.com/dsb-lab/qlivecell) in combination with the StarDist 2D pretrained model, **2d_versatile_fluo** (CITE). Segmentation masks were classified as cells based on the following criteria: a) They are present in at least two z-planes. b) They exceed a size threshold in their center plane. The center plane was defined as the z-plane with the highest summed fluorescence intensity within the segmented mask. The 2D center of the mask was calculated as the centroid of this plane. This 3D coordinate was used consistently as the cell center, as crosstalk between channels within the segmented masks increases with distance from the center plane.

The size threshold for the masks was determined from the distribution of mask areas across all segmented objects in their center plane. Our segmentation pipeline was applied to 96-hour gastruloids. To define the threshold, we fitted a kernel density estimation to the histogram of mask areas, using the smallest kernel width that produced a single local minimum. This local minimum (33.34 ± 2.8 µm²) was used to distinguish between segmented cells and segmented cellular debris. The cell count for each population is obtained as the number of segmented objects in the correspondent channel to that population after debris removal. The number of segmented objects removed are used for the debris quantification.

### Classification and quantification of apoptotic events

Apoptotic events are quantified using the segmentation of the Caspase-3 immunostaining. Due to the high variability in the Caspase-3 signal of apoptotic cells, we manually curated the segmentation results to ensure accuracy using the QLIVECELL software. We divided our analysis into three apoptotic stages, based on a combination of nuclear morphology and the Caspase-3 staining pattern: 1) **Early apoptosis**: The nucleus appears intact, with surrounding cytoplasmic Caspase-3 staining. 2) **Mid apoptosis**: The nuclear membrane begins to dissolve, and the

Caspase-3 signal mixes with the nuclear signal. Chromatin granules form, creating chromatin-free regions between them, resulting in a more intense and fragmented nuclear signal. 3) **Late apoptosis**: The nuclear size is reduced, and chromatin is fully compacted. Our classification aligns with stages described in CITE: early apoptosis corresponds to stages 1 and 2, mid apoptosis to stages 3 and 4, and late apoptosis to stage 5. For segmenting late apoptotic cells, we applied intensity and size thresholds to ensure the identification of debris. The size threshold was established using the same approach as for debris removal, but in this case, we excluded larger objects and retained smaller particles. The intensity threshold was based on the mean Caspase-3 signal of early apoptotic events, retaining only those with high signal intensity.

### Radial distribution of cells

Spatial distribution of cells in the gastruloids is computed using their center-edge relative position. The relative position of each cell is computed as the distance to the 3D centroid of the gastruloid divided by the sum of the distance to that centroid and the distance to the closest gastruloid edge. The edge of the gastruloid for each z-plane is obtained using the scikit-image implementation of morphological active contours without edges CITE.

### Cell density

We compute the local cell density ρlocal using a nearest neighbors approach. Specifically, we picked the 10 closest nearest neighbors for each cell using the K-nearest neighbor algorithm (CITE), and defined the local density as the cubic mean distance to the neighbors. Cells that are closer than 0.25*cell diameters are discarded from the neighborhood.

### Neighborhood composition

Neighborhood composition is defined as the fraction of cells belonging to a specific population within the neighborhood of a given cell. The neighborhood is determined using the k-nearest neighbors algorithm (CITE). Cells that are closer than 0.25*cell diameters are discarded from the neighborhood.

### Transcriptomic data preprocessing

The quality of the raw sequencing files was assessed using FastQC (version 0.11.9) (Andrews, 2010). Adapter trimming and low-quality filtering were conducted with TrimGalore! (version 0.6.7) (Felix Krueger, https://github.com/FelixKrueger/TrimGalore), removing reads with a Phred score below 20. Post-trimming, the cleaned reads were re-evaluated with FastQC to verify the effectiveness of the trimming and filtering processes.

The processed sequencing data were aligned to the mouse genome (GRCm38.p6) using Gencode annotation (version 25) and the STAR aligner (version 2.7.11b)^51^. Alignments were performed with default settings and uniquely mapped read counts were generated using the --quantMode option in STAR. A total of 3,695,039,036 uniquely mapped reads were obtained, and these served as input for downstream analyses and visualization. The final count matrix was created by concatenating the assembled reads in a Python 3 environment.

### Downstream analysis and visualization

All downstream statistical analyses and visualizations were conducted in a Python 3.12.8 environment. The median ratio normalization was computed using the PyDESeq2 (version 0.4.0) module, and the normalized counts were log1p-transformed for further analysis.

For dimensionality reduction, the scikit-learn library (version 1.2.2) was employed. The most variant genes (MVG) were selected, which were then used as input for principal component analysis (PCA).

To map bulk samples to embryonic developmental stages, the Euclidean distance was calculated between the averaged expression conditions and pseudobulk embryo conditions from the Pijuan-Sala dataset (E6.5–E8.5) {Pijuan-Sala, 2019 #2707}. This analysis facilitated temporal alignment of the data to embryonic developmental stages. A similar approach was used to align samples to spatial information from the same embryonic dataset (Anterior-Posterior axis) at E8.5.

For visualization purposes, modules seaborn (version 0.13.2) and matplotlib (version 3.9.2) were utilised for plotting.

The complete code is available on GitHub at https://github.com/stembryo-lab/cell_competition_gastruloids

### Signaling modulators, inhibitors, and ROS scavenger treatments

Gastruloids were either treated from 48-72h after aggregation with a pulse of CHIR (potent Wnt/β- catenin signaling activator) (Chi99021, Sigma Cat#SML1046-5MG) at 3 μM, or received alternative treatments in the absence of ChIR. Alternative treatment regimens were conducted as follows: 48h-72h Activin A (100 ng/mL, QKINE #Qk005), which mimics Nodal signaling; 48h-120h IWP-2 (pan-inhibitor of Wnt ligand secretion, 1 μM, Sigma #I0536); 48h-120h XAV-939 (inhibitor of the Wnt canonical pathway, 1 μM, Tocris #3748); 48h-120h SB431542 (Activin/Nodal signaling inhibitor, 10 μM, Tocris #1614); 48h-120h BMP4 (1ng/mL, R&D #5020-BP); 48h-120h LDN193189 (BMP signaling inhibitor, 100 nM, Tocris #6053); 48h-120h Fgf2 (12.5ng/mL, Peprotech #450-33); 48h-120h PD0325901 (Fgf signaling inhibitor, 1 μM, Sigma #Pz0162).

The reactive oxygen species (ROS) scavenger N-acetyl-L-cysteine (NAC, Sigma #A9165) was added to gastruloids at 20 μM, 100 μM, 500 μM, 2500 μM, or 12500 μM in addition to the standard ChIR treatment from 48h to 72h.

### Heterochronic gastruloids

Pure gastruloids were generated from 300 WT-emiRFP cells, 300 WT-mCherry cells, or 75 p53KO-emiRFP cells per well as described above at different days and left to grow until indicated time points. E.g. to obtain cells from 24h or 72h gastruloids, pure gastruloids were generated 1 or 3 days before the assembly, while WT-mCherry cells were always taken from 48h gastruloids generated 2 days before the assembly. On the day of heterochronic assembly, gastruloids were harvested and pooled within the same condition. Each set of pooled gastruloids was rinsed once in warm PBS -/- and incubated for 5 minutes in Accutase cell detachment solution (LabClinics #ACC-1B) at 37°C, agitating the tubes regularly. For each condition, a single cell suspension was then obtained by pipetting up and down ∼15 times. Single cell suspension was then centrifuged for 3 minutes at 200 xg. The supernatant was carefully discarded, and cells were counted in 100-250 μL N2B27. Different heterochronic conditions were then generated by mixing 48h WT-mCherry gastruloid cells with WT-emiRFP or p53KO-emiRFP cells from gastruloid of different timepoints in CHIR 3μM or in DMSO. Each of the master mixes was designed to allow seeding a mix of 1250 WT-mCherry cells with 1050 WT-emiRFP or 480 p53KO-emiRFP cells per 150μL in each well of a 96-well plate, which are the numbers observed in respective mosaic gastruloids at 48h, with extra cells to account for cell loss. The cell mixes were seeded into each well of a 96-well plate in 150 μL volume, the plates were centrifuged for 1min at 150 xg and placed back in the incubator. Every 24h until the WT-mCHerry reference cells had reached 120h (3 days total), fresh N2B27 was added to the gastruloids.

### Epi gastruloids

The protocol for generating Epi-gastruloids was originally established in the doctoral thesis of Dr. Shlomit Edri (Edri 2019a; Doctoral Thesis). mESC in ES/LIF medium were seeded onto 6 wells coated with fibronectin (2 µg/ cm^2^) (Bio-Techne, Cat#1918-FN-02M) or vitronectin (0.5 µg/ cm^2^) (Fisher Scientific, Cat#A14700). The following day, the medium was exchanged for N2B27 (NDiff227-custom, Takara, Cat# XA0530) supplemented with 12 ng/mL bFGF2 (Preprotech, Cat#450-33 100µg) + 25 ng/mL Activin A (QKine, Cat#Qk005-100ug). The following day, cells were gently lifted off the well as clumps using a 0.5 mM EDTA PBS-/- solution and passaged onto a new fibronectin or vitronectin coated well in N2B27+AA+bFGF2. After the passage 2, cell colonies start appearing flatter with spread out cell filopodia at the edge. On the 3^rd^ passage, cells reached an Epi-like state and were further converted to EpiCE cells. For this, cells were enzymatically dissociated to single cells using Accutase and seeded as 5*10^4^ cells per 6-well in N2B27+AA+FGF supplemented with 5 µM ROCK inhibitors (Y-27632, Selleckchem, Cat#S1049-10MG). On the second day, the medium was exchanged to N2B27+ 20ng/mL bFGF2 without Activin A. On the third day, the medium was exchanged to N2B27+ 20ng/mL bFGF2 supplemented with 3 µM ChIR (Chi99021, Sigma Cat#SML1046-5MG). 24h after the ChIR addition, these cells are referred to as EpiCE. EpiCE cells were enzymatically dissociated to single cells using Accutase. After a washing step, cells were resuspended in EpiGastruloid medium (N2B27 + 20ng bFGF + 3 µM ChIR) and counted. As a default, EpiGastruloids were seeded as 600 total cells in 40 µL medium per U-bottom 96-well (Ultra-low attachment plates Grainer Cat#650970). WT+WT mosaic EpiGastruloids were seeded as 300 WT-mCherry cells + 300 WT-emiRFP670 cells. WT+KO mosaic EpiGastruloids were seeded as 300 WT-mCherry cells + 50 p53KO-emiRFP670 cells. 24h after EpiGastruloid aggregation, 150 µL fresh EpiGastruloid medium was added to each well. Every following day, 150 µL medium were removed and replaced with fresh EpiGastruloid medium.

---

**Extended Data Figure 1.**
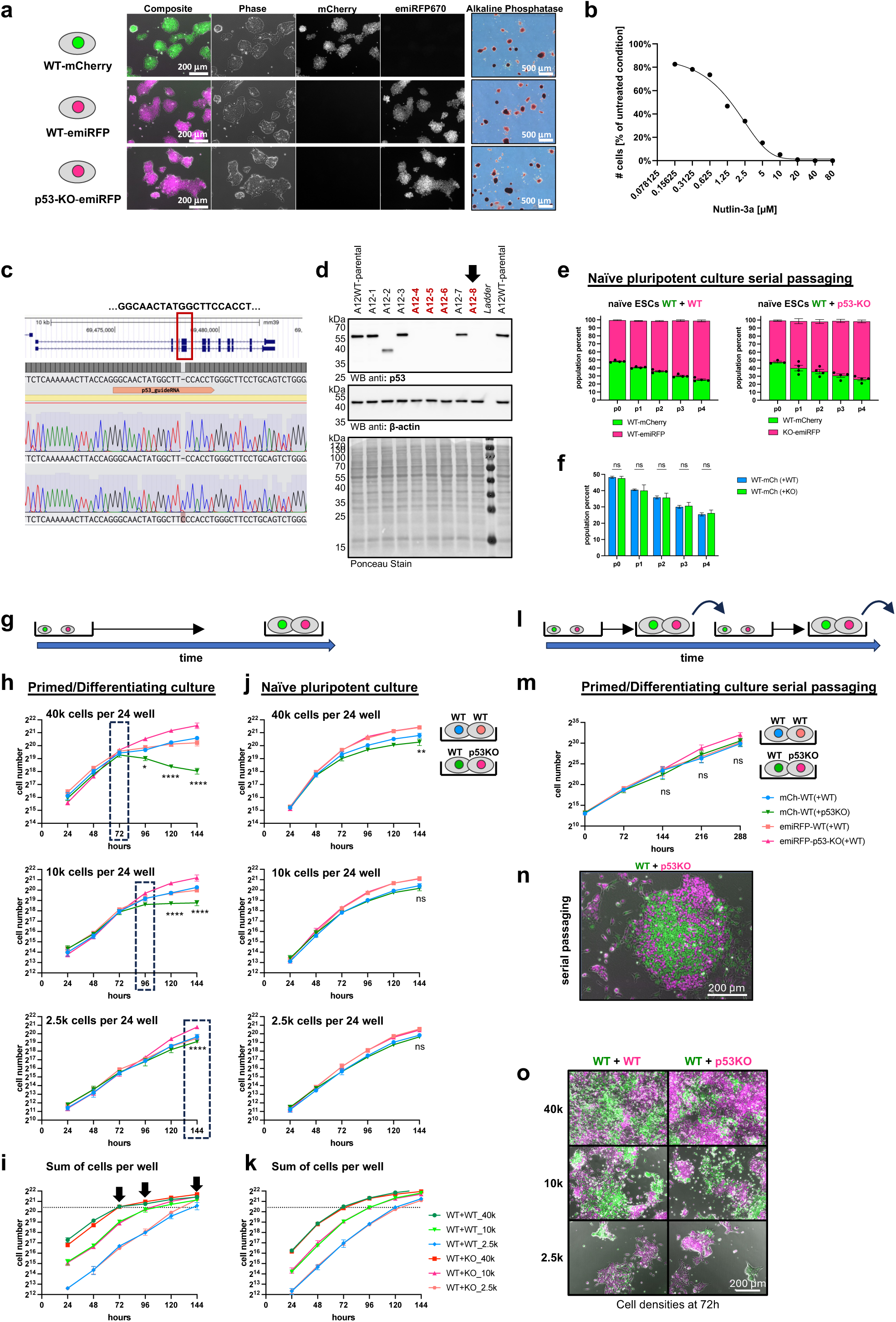
2D cell competition occurs only at confluence during differentiation. **a**, Overview of H2B-mCherry and H2B-emiRFP670 tagged mESC clones used in the study. Widefield fluorescent micrographs of adherent maintenance cultures, and brightfield image of alkaline phosphatase staining (right panels). Scale bar denotes 200 µm and 500 µm, respectively. **b**, Selection kill curve depicting WT mESC viability after 96h treatment with varying Nutlin-3a concentrations, expressed as percent viable cells of untreated control. **c**, Sequencing alignment of WT cells and p53KO-emiRFP clone A12-8 to *Trp53* reference gene. Frame shifting +1 insertion in p53KO clone. **d**, Western blot analysis of WT-emiRFP parental clone (“A12-WT”) and daughter clones stained against p53 (upper panel) and loading control β-actin (middle panel) confirms loss of p53 protein in clone A12-8, which is used as “p53KO-emiRFP” clone in this study. Ponceau stained membrane functions as quality control of protein transfer (lower panel). **e**, Flow cytometric analysis of population composition of mixed naïve pluripotent WT+WT or WT+p53KO co-cultures. Bar graphs depict percentage of mCherry+ (green) and emiRFP+ (magenta) cells from passages p0 to p4. n=4 independent experiments. **f**, Comparison of WT-mCherry population percentage from mixed co-cultures with WT-emiRFP or p53KO-emiRFP cells depicted in panel **e**. n=4 independent experiments. ns, not significant. **g**, Schematic of co-culture experiments under pluripotent or differentiating conditions without passaging. **h**, 2D cell competition experiment under differentiating conditions (N2B27+AA+FGF2). Growth curves of WT-mCherry (blue line) cells co-cultured with WT-emiRFP (orange line) cells are overlaid with WT-mCherry (green line) cells co-cultured with p53KO-emiRFP (magenta line) cells. 1:1 mixed cell populations were seeded as 40,000 (top), 10,000 (middle), or 2,500 (bottom) total cells per 24 well. Dashed boxes indicated onset of cell competition. n=3 independent experiments. Statistical comparison of WT-mCherry cells from WT+WT or WT+p53KO condition shown. *, p<0.05; ****, p<0.0001 **i**, Sum of total cells per well from experiments depicted in panel **h**. Arrows demarcate the timepoints of onset of competition in panel **h**. Color scheme follows legend of panel **k**. **j**, 2D cell competition experiment under naïve pluripotent culture conditions. Growth curves of WT-mCherry (blue line) cells co-cultured with WT-emiRFP (orange line) cells are overlaid with WT-mCherry (green line) cells co-cultured with p53KO-emiRFP (magenta line) cells. 1:1 mixed cell populations were seeded as 40,000 (top), 10,000 (middle), or 2,500 (bottom) total cells per 24 well. n=3 independent experiments. Statistical comparison of WT-mCherry cells from WT+WT or WT+p53KO condition shown. ns, not significant; **, p<0.01 **k**, Sum of total cells per well from experiments depicted in panel **k**. **l**, Schematic of co-culture experiments under differentiating conditions with continuous passaging. **m**, 2D cell competition experiment under differentiating conditions (N2B27+AA+FGF2) with passaging every 72h. Growth curves of WT-mCherry (blue line) cells cocultured with WT-emiRFP (orange line) cells are overlaid with WT-mCherry (green line) cells cocultured with p53KO-emiRFP (magenta line) cells. n=3 independent experiments. Statistical comparison of WT-mCherry cells from WT+WT or WT+p53KO condition shown; ns, not significant. **n**, Widefield fluorescent micrograph of a representative mixed colony during the second passage of differentiating co-cultures depicted in panel **m**. **o**, Representative widefield fluorescent micrographs of mixed co-cultures after 72h in differentiating conditions quantified in panel **h**. All values depicted as mean ±SEM throughout this figure.

**Extended Data Figure 2.**
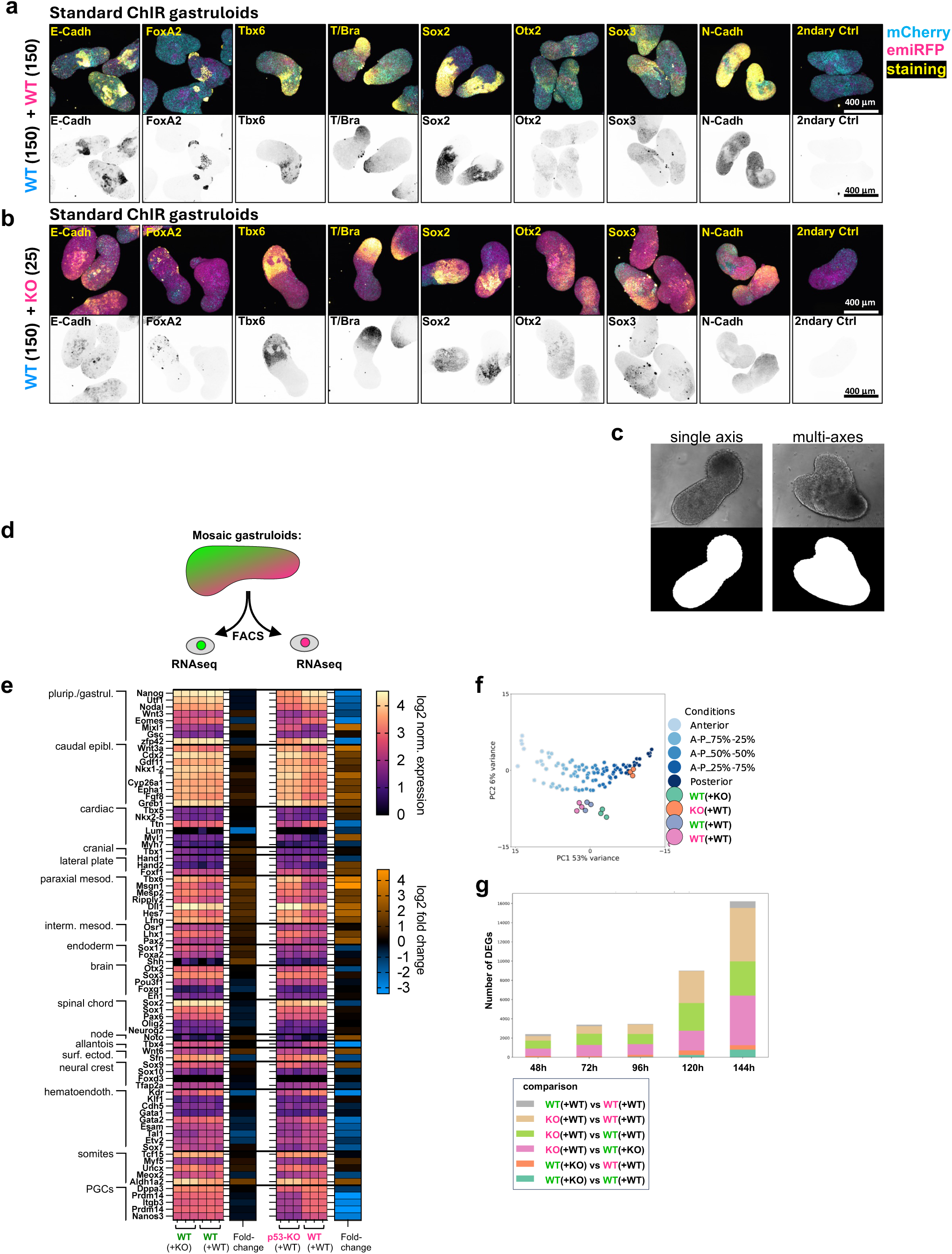
Characterization of mosaic gastruloid fate and morphology. **a-b**, Fate marker staining of mosaic WT+WT (**a**) and WT+p53KO (**b**) gastruloids at 120h. Maximum intensity projections of spinning disk confocal imaged, fixed, and immunostained gastruloids. Micrographs depicted as composite (top panel) of mCherry (blue), emiRFP (magenta), and marker staining (yellow), or inverted marker staining alone (lower panel). Scale bar denotes 400 µm. **c**, Example phase contrast micrographs of single-axis and multi-axes gastruloids at 120h and respective masks used for analysis of morphological features. **d**, Schematic of FACS isolated mCherry+ and emiRFP+ cells from mosaic gastruloids prior to RNA sequencing analysis. **e**, Heatmap of developmental lineage marker expression profile from mRNA bulk sequencing analysis of pre-sorted mosaic gastruloids following schematic of panel **d**. Log2 normalized transcript levels of biological triplicates are depicted as magma heatmap. Log2 fold-change of WT-mCherry cells from WT+WT vs from WT+p53KO gastruloids, and log2 fold-change of p53KO-emiRFP vs WT-emiRFP cells from mosaic gastruloids depicted as blue to orange heatmap adjacent. **f**, Principal component analysis of clonal populations sorted from 120h mosaic gastruloids together with anterior-to-posterior pseudobulk subsets of embryonic stage E8.5 transcriptomes (blue shades). **g**, Number of significant differentially expressed genes (DEG) between pairs of FACS-isolated clonal populations from mosaic gastruloids at varying timepoints. All values depicted as mean ±SEM throughout this figure.

**Extended Data Figure 3.**
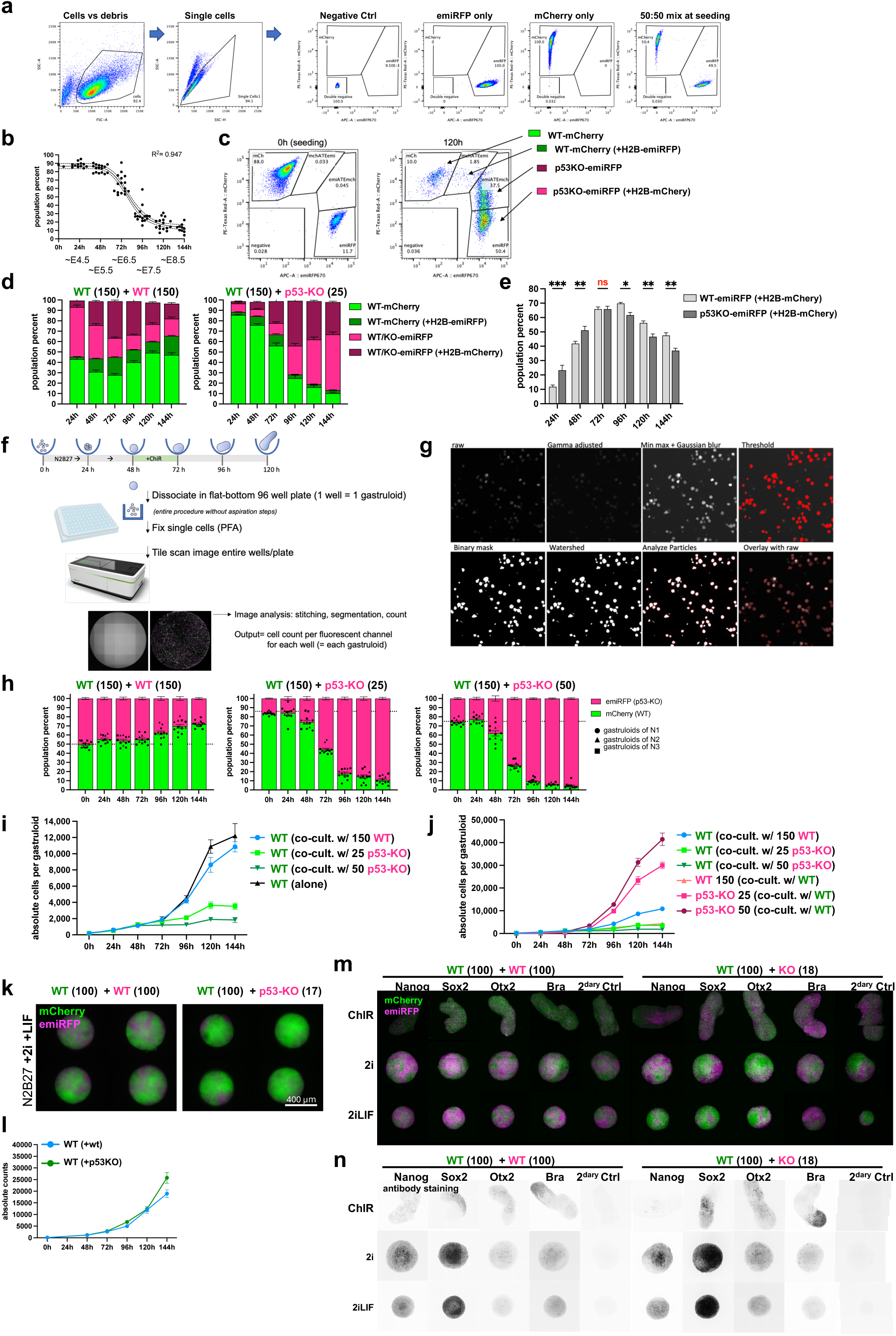
Phagocytosis, growth curve, and pluripotent aggregate analysis. **a,** Gating strategy of flow cytometry analysis of mCherry and emiRFP mixed populations. (corresponds to Fig. 2h-i) b, WT-mCherry population percentages in WT-p53KO mosaic gastruloids measured by single gastruloid flow cytometry (corresponds to Fig. 2i). Only mCherry WT datapoints are depicted together with fitted logistic decline curve, R^2^=0.947. n=12-16 gastruloids from 3 independent experiments. c, Flow cytometric gating strategy for analysis of neighbor phagocytosis as determined by double positive mCherry+/emiRFP+ events. d, Flow cytometric analysis of neighbor phagocytosis in WT+WT (left) or WT+p53KO (right) mosaic gastruloids at varying timepoints. Light colors indicate cells that have not taken up fluorescent H2B-fluorophore material from respective other neighbors, dark colors indicate cells that have taken up foreign fluorescent material from neighbors. n=12-16 gastruloids from 3 independent experiments. e, Comparison of percentages of WT-emiRFP and p53KO-emiRFP cells that have phagocytosed WT-mCherry neighboring cell debris in mosaic gastruloids at varying timepoints. n=12-16 gastruloids from 3 independent experiments. Statistical analysis between conditions within same timepoints depicted; *, p<0.05; **, p<0.01; ***, p<0.001; ns, not significant. f, Schematic of experimental workflow of determining absolute cell numbers per fluorescent population per gastruloid by dissociation, imaging, and nuclear segmentation. g, Overview of image processing steps during nuclear segmentation of 2D tiled images of individually dissociated gastruloids. h, Population percentages of mCherry and emiRFP populations in mosaic gastruloids as determined from absolute cell number counts with methodology outlined in panel f. n=12 gastruloids from 3 independent experiments. **i-j**, Absolute growth curves of WT-mCherry cells (**i**) in mosaic gastruloids seeded together with 150 WT-emiRFP (blue), 25 p53KO-emiRFP (light green), 50 p53KO-emiRFP (dark green) cells, or alone (black line), or WT-emiRFP and p53KO-emiRFP cells (**j**) from the same mosaic gastruloids. n=12 gastruloids per datapoint from 3 independent experiments. (corresponds to Fig. 2k). **k**, Representative widefield fluorescent images of pluripotent mosaic 3D aggregates grown in N2B27+2i+LIF. Mosaic aggregates of WT+WT (left) or WT+P53KO (right) gastruloids imaged at 120h after aggregation. Scale bar denotes 400 µm. (corresponds to condition without LIF in Figure 2l). **l**, Absolute growth curves of WT-mCherry cells in mosaic pluripotent aggregates seeded together with WT-emiRFP (blue) or p53KO-emiRFP (dark green) cells. n=12 gastruloids per datapoint from 3 independent experiments. **m-n**, Fate marker staining of mosaic WT+WT (left) and WT+p53KO (right) aggregates at 120h grown following the standard gastruloid protocol or remaining as pluripotent aggregates in 2i or 2i+LIF. Maximum intensity projections of confocal laser scanning micrographs of fixed, and immunostained gastruloids. Micrographs depicted as composite (**m**) of mCherry (green) and emiRFP (magenta), or as inverted marker staining alone (**n**). All values depicted as mean ±SEM throughout this figure.

**Extended Data Figure 4.**
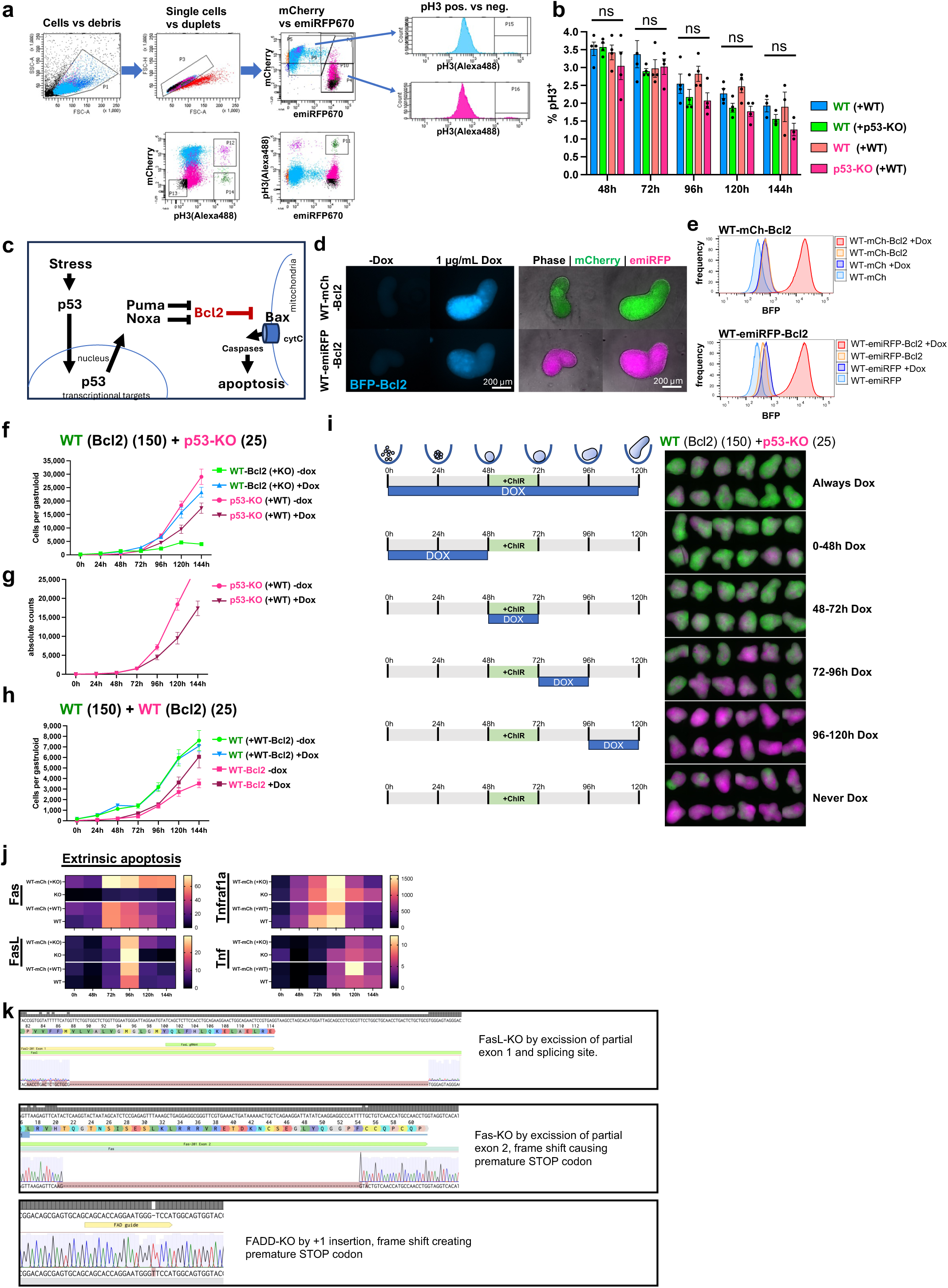
Cell competition in gastruloids is proliferation and extrinsic apoptosis independent. **a**, Flow cytometry gating strategy to quantify phospho-histone3 (pH3) positive percentage of mCherry+ or emiRFP+ cells within mosaic gastruloids. **b**, Flow cytometric quantification of pH3 positive percentage of different cell populations within WT+WT or WT+p53KO mosaic gastruloids at varying time points. n=4 independent experiments. Two-Way ANOVA with post-hoc Tukey’s multiple testing within each timepoint; ns, not significant. n=4 independent experiments. **c**, Schematic of stress response pathway downstream of p53, highlighting the dominant negative role of anti-apoptotic Bcl2. **d**, Fluorescent micrographs demonstrating doxycycline-inducible nature of Bcl2 overexpression clones, as detected by co-expressed BFP. Scale bar denotes 400 µm. **e**, Flow cytometry histograms validate the doxycycline-inducible nature of Bcl2 overexpression clones, as detected by co-expressed BFP fluorophore. **f-g**, Overlay of absolute growth curves from Fig. 2c and Fig. 2d of all cell populations (**f**), or only of p53KO-emiRFP (**g**) in response to Bcl2 induction in WT-mCherry-Bcl2 cells. **h**, Overlay of absolute growth curves from Fig. 2f and Fig. 2g of all cell populations. **i**, Schematic of temporal Bcl2 induction series in WT-mCherry-Bcl2 clones in mosaic WT+p53KO gastruloids, together with representative fluorescent montages of resulting 120h gastruloids from corresponding temporal induction protocols. **j**, Heatmap of temporal gene expression level dynamics of extrinsic apoptosis pathway genes. FACS-isolated cell populations from mosaic gastruloids depicted separately in rows and timepoints from 48h-144h after aggregation in columns. Each datapoint is a mean of 3 biological replicates. **k**, Sequencing validation of FasL-, Fas-, and Fadd-knockout clones. All values depicted as mean ±SEM throughout this figure.

**Extended Data Figure 5.**
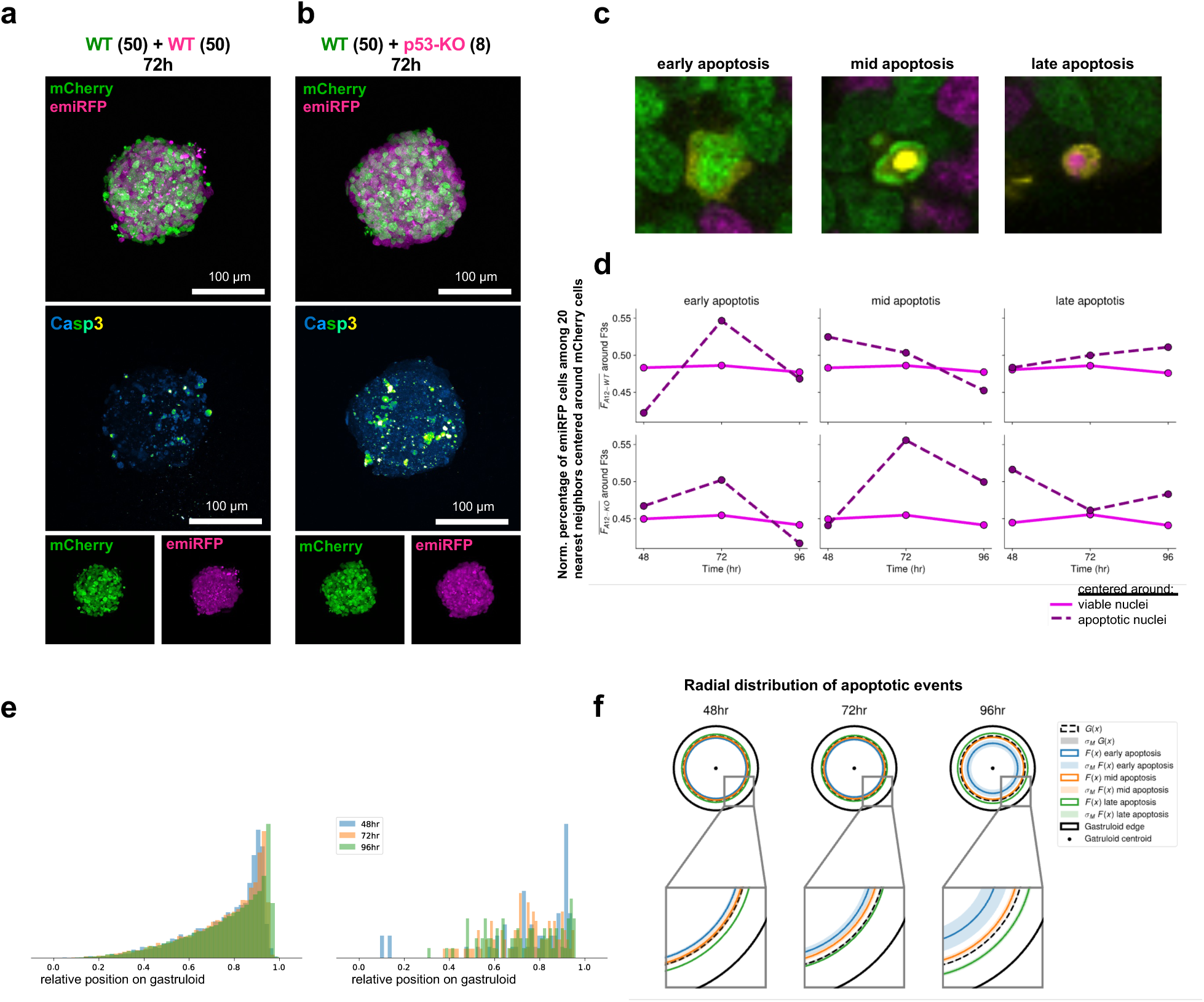
Apoptotic events are biased toward p53KO neighborhoods and inner regions of gastruloids. **a-c**, Representative maximum intensity projection of confocal fluorescent micrographs depicting mosaic WT+WT (**a**) or WT+p53KO (**b**) gastruloids at 72h after aggregation. Caspase 3 (Casp3) immunofluorescent staining depicted in separate panels as GreenFireBlue heatmap (middle panels). **c**, Example cropped fluorescent micrographs of early apoptotic, mid apoptotic, and late apoptotic events as identified by caspase 3 staining (yellow) and nuclear morphology. **d**, Neighborhood analysis of segmented nuclei. Normalized percentage of emiRFP+ nuclei among 20 nearest neighbors centered around mCherry+ nuclei. Solid lines depict neighborhoods around viable cell nuclei. Dotted lines represent neighborhoods centered around apoptotic events. **e**, Histogram of total quantified viable (left) and apoptotic (right) nuclei along center-to-edge axis as measured within an expanding shell within 3D segmented gastruloids. **f**, Radial distribution of all nuclei (G_(x)_, black dotted ring), early (blue), mid (orange) or late (green) apoptotic events within gastruloids at 48h, 72h, and 96h after aggregation.

**Extended Data Figure 6.**
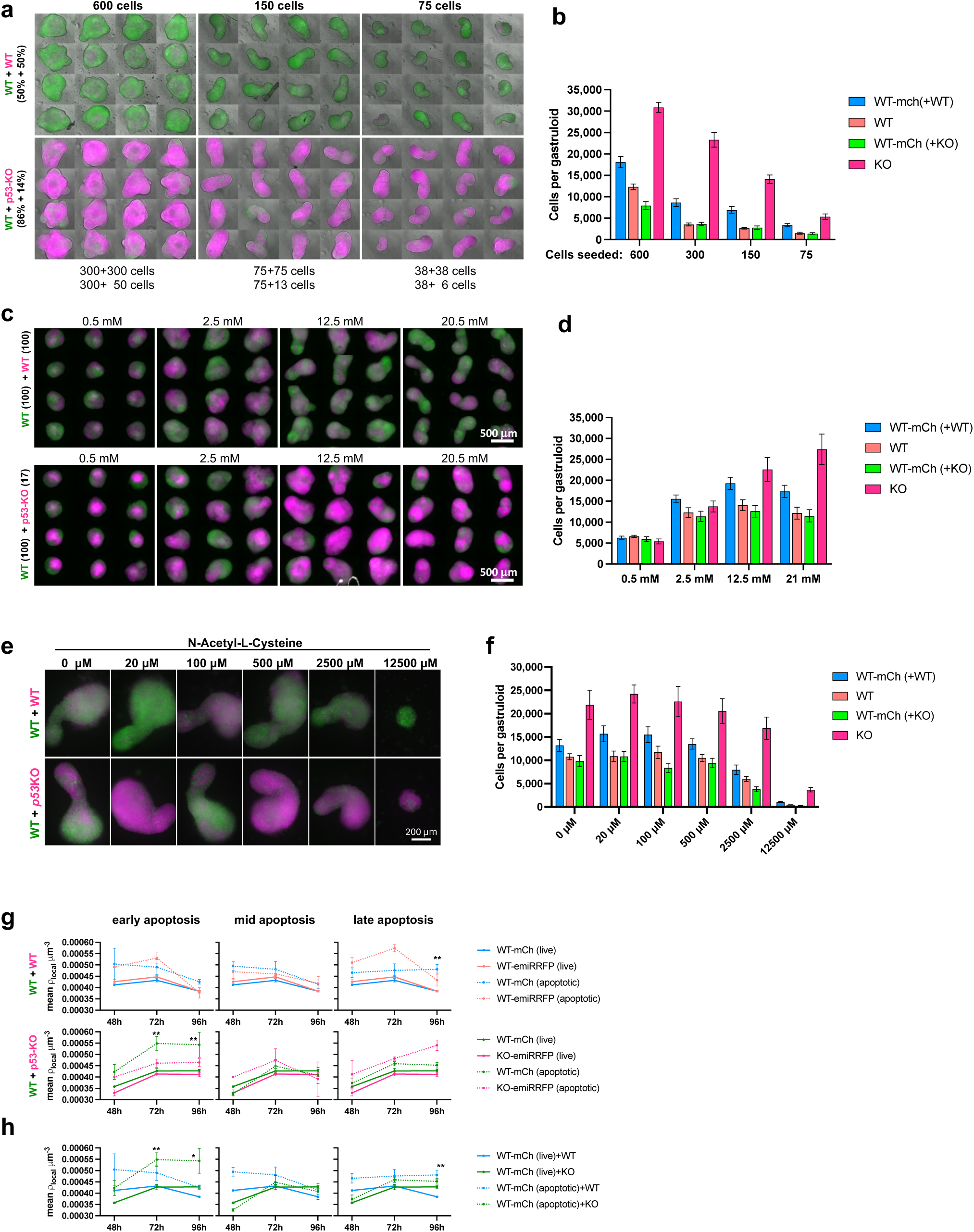
Size, glucose, ROS-scavenger titration and local density analysis. **a,** Fluorescent widefield micrographs of 120h mosaic WT+WT (top) or WT+p53KO (bottom) gastruloids seeded from 600, 150, or 75 total cells. **b,** Absolute cell counts quantification of individual populations at 120h in mosaic gastruloids as depicted in panel a. n=12 gastruloids from 3 independent experiments. **c,** Fluorescent widefield micrographs of 120h mosaic WT+WT (top) or WT+p53KO (bottom) gastruloids grown in varying concentrations of glucose. Scale bar denotes 500 µm. **d,** Absolute cell counts quantification of individual populations at 120h in mosaic gastruloids as depicted in panel c. n=24 gastruloids from 4 independent experiments. **e**, Representative fluorescent widefield micrographs of 120h mosaic WT+WT (top) or WT+p53KO (bottom) gastruloids grown in varying concentrations of the ROS scavenger N-acetyl-L-cysteine. Scale bar denotes 200 µm. **f**, Absolute cell count quantification of individual populations at 120h in mosaic gastruloids as depicted in panel e. n=11 gastruloids from 3 independent experiments. **g**, Mean local density in mosaic WT+WT (top) or WT+p53KO (bottom) gastruloids at 48h, 72h, or 96h as determined by the distance between 3D segmented nuclei. Analysis centered around viable nuclei (solid lines) or early-, mid-, or late apoptotic events (dashed lines). Line color indicates which clone-specific nuclei the analysis was centered around. WT-mCherry (+WT), blue; WT-mCherry (+KO), green; WT-emiRFP (+WT), orange; p53KO-emiRFP (+WT), magenta. Mean of all nuclei from 3-5 gastruloids per condition. Only statistics between WT-mCherry in WT+WT vs WT+p53KO gastruloids from the same condition are depicted; *, p<0.05; **, p<0.01. **h**, Overlaid WT-mCherry graphs from WT+WT (blue) and WT+p53KO (green) graphs of panel g. *, p<0.05; **, p<0.01. All values depicted as mean ±SEM throughout this figure.

**Extended Data Figure 7.**
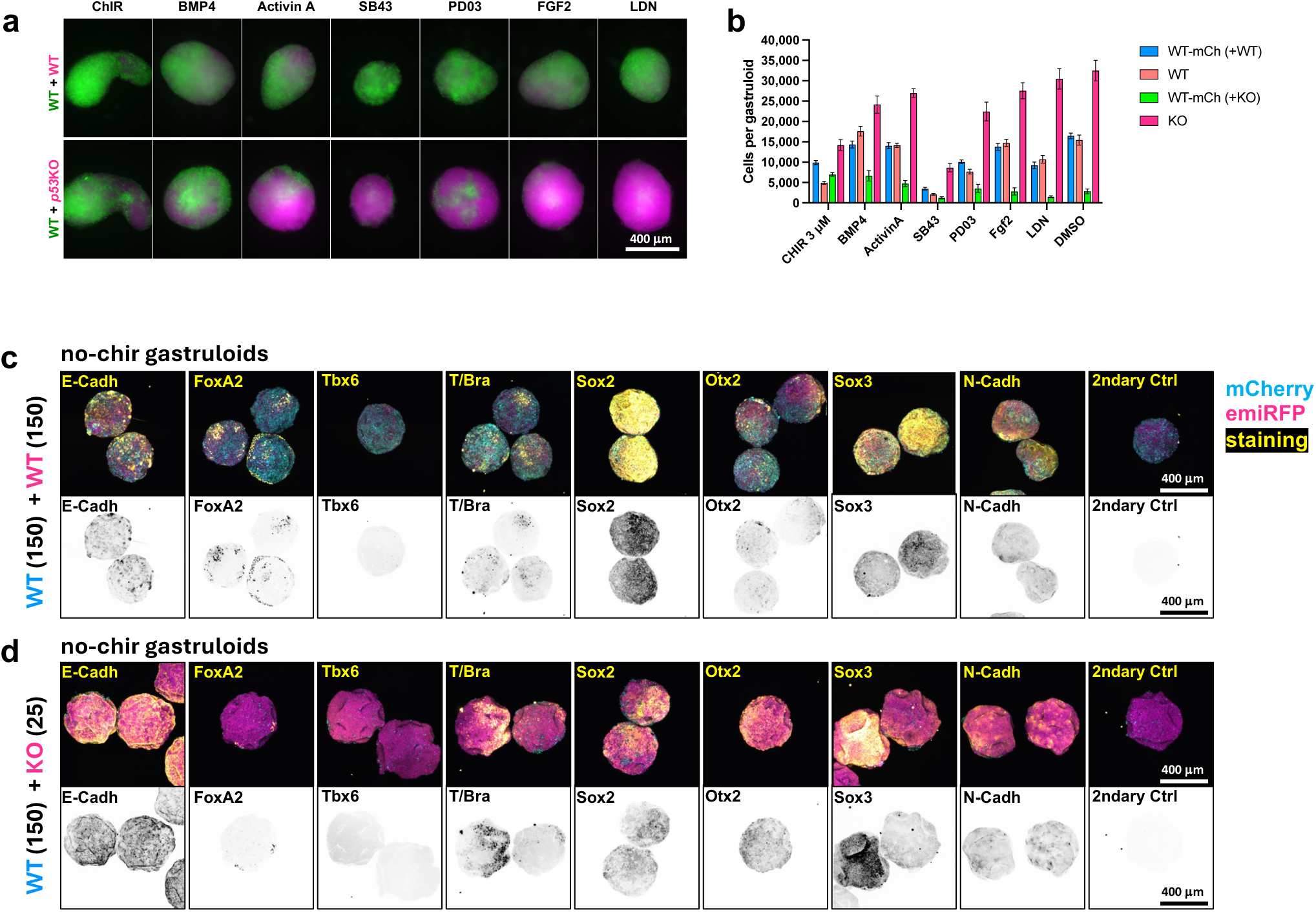
Effects of signalling perturbations on cell competition and fate specification. **a-b**, Representative fluorescent micrographs (**a**) of 120h mosaic WT+WT (top) and WT+p53KO (bottom) gastruloids treated with various developmental signal modulators, and absolute counts of different cell populations at 120h (**b**). Gastruloids were treated with 3 µM ChIR (48-72h), 1ng/mL BMP4 (48-120h), 100ng/mL Activin A (48-72h), 10 µM SB431542 (48-120h), 1 µM PDO325901, 12.5 ng/mL FGF2 (48-120h), 100 nM LDN193189 (48-120h), or DMSO carrier control. n=12 gastruloids from 3 independent experiments. **c**, Fate marker staining of mosaic WT+WT (**a**) and WT+p53KO (**b**) gastruloids at 120h grown without ChIR treatment (DMSO control). Maximum intensity projections of spinning disk confocal imaged, fixed, and immunostained gastruloids. Micrographs depicted as composite (top panel) of mCherry (blue), emiRFP (magenta), and marker staining (yellow), or inverted marker staining alone (lower panel). All values depicted as mean ±SEM throughout this figure. All scale bars denote 400 µm.

**Extended Data Figure 8.**
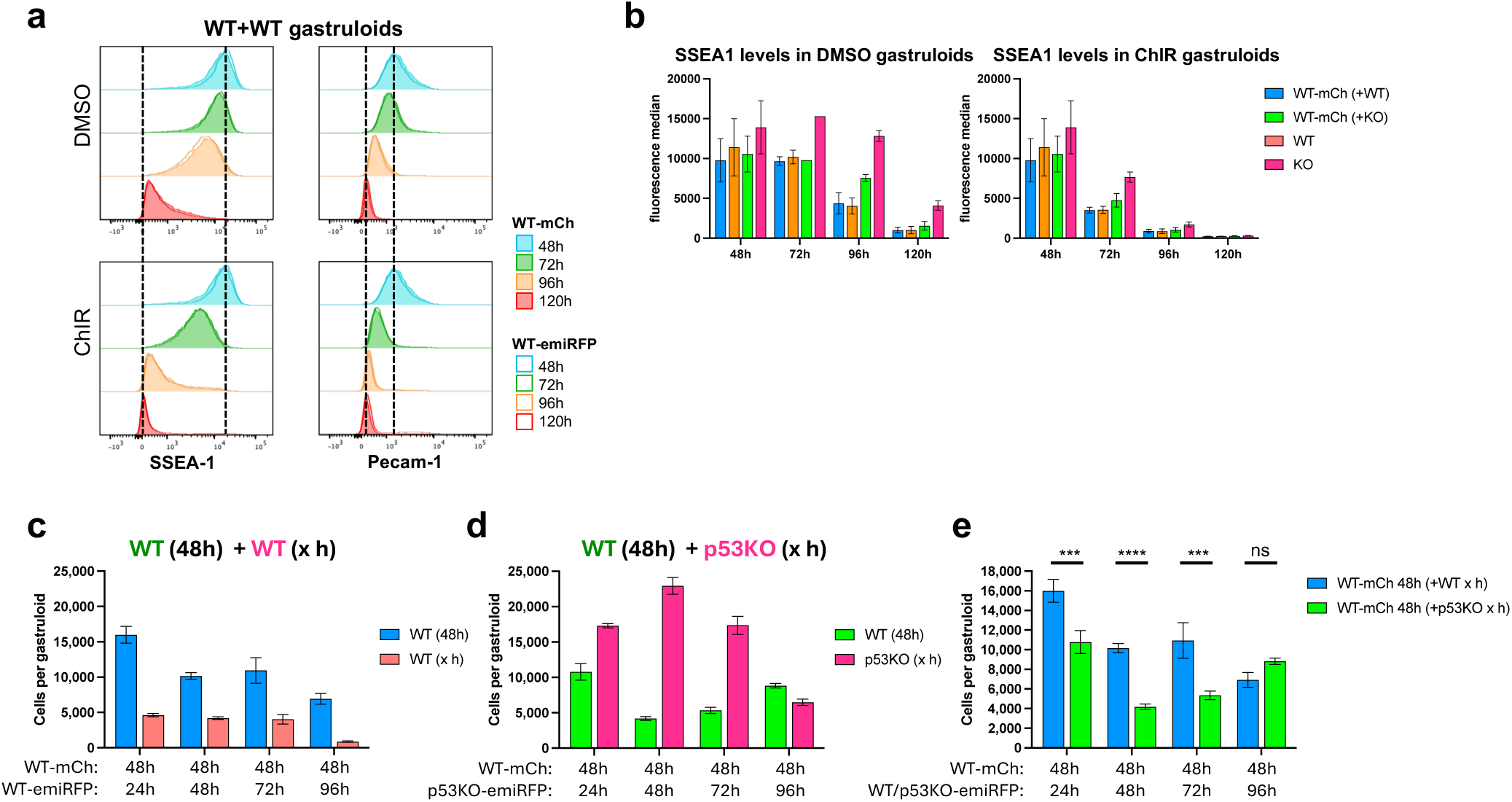
Temporal analysis of stemness marker downregulation and heterochronic gastruloids. **a**, Flow cytometric quantification of SSEA1 and Pecam-1 pluripotency surface marker expression in WT-mCherry (filled histograms) and WT-emiRFP (open lines) from mosaic gastruloids treated with ChIR (lower panels) or DMSO control (upper panels) analyzed at 48h, 72h, 96h, or 120h. **b**, Quantification of SSEA-1 surface marker expression in ChIR or DMSO control treated mosaic gastruloids at 48h, 72h, 96h, or 120h after aggregation, as determined by flow cytometry depicted in panel **a** and Fig. 6c. n=3 independent experiments. **c-d**, Heterochronic mosaic gastruloids created from 48h WT-mCherry gastruloid cells with WT-emiRFP or p53KO-emiRFP cells from gastruloid of varying timepoints. Absolute counts of cell populations in mosaic WT+WT (**c**) or WT+p53KO (**d**) gastruloids at 120h, 72h after reaggregation, depicted. Heterochronic experiments in this panel conducted with home-made N2B27, while heterochronic experiments in Fig. 6 were conducted with commercial N2B27. n=12 gastruloids from 3 independent experiments. **e**, Comparative overlay graph of WT-mCherry populations from panels **c** and **d**. Only statistics between WT-mCherry cell numbers in WT+WT vs WT+p53KO gastruloids of the same heterochronic condition depicted; ***, p<0.001; ****, p<0.0001; ns, not significant. All values depicted as mean ±SEM throughout this figure.

**Extended Data Figure 9.**
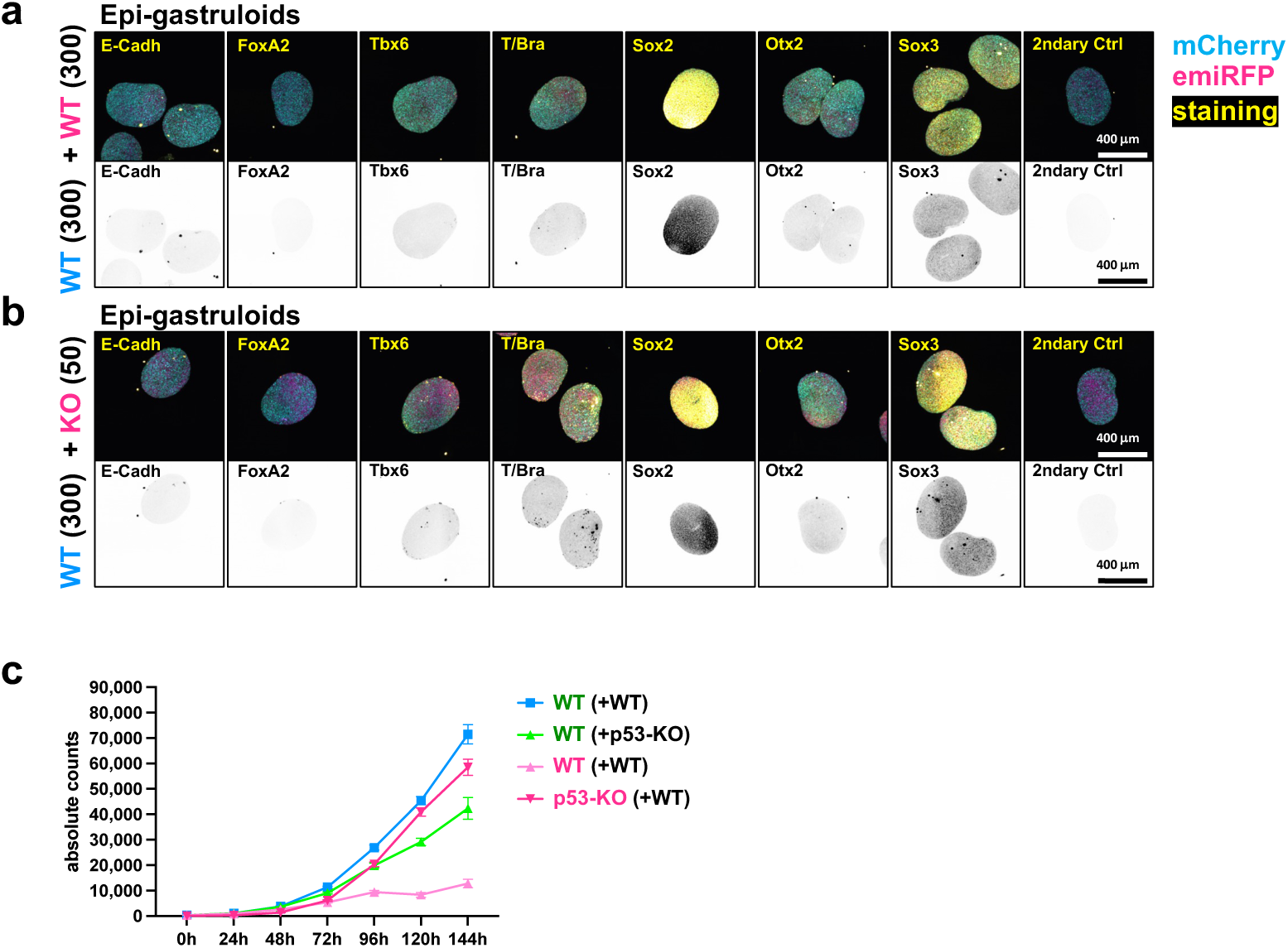
Analysis of EpiGastruloid cell fates. **a-b**, Fate marker staining of mosaic WT+WT (**a**) and WT+p53KO (**b**) EpiGastruloids at 72h. Maximum intensity projections of spinning disk confocal imaged, fixed, and immunostained EpiGastruloids. Micrographs depicted as composite (top panel) of mCherry (blue), emiRFP (magenta), and marker staining (yellow), or inverted marker staining alone (lower panel). Scale bars denote 400 µm. **c**, Absolute growth curves of all cell populations in mosaic EpiGastruloids. Corresponding to Fig. 7i. n=17 gastruloids per datapoint from 3 independent experiments.

